# Hao-Fountain syndrome protein USP7 controls neuronal differentiation via BCOR-ncPRC1.1

**DOI:** 10.1101/2024.08.04.606521

**Authors:** Joyce Wolf van der Meer, Axelle Larue, Jan A. van der Knaap, Gillian E. Chalkley, Ayestha Sijm, Leila Beikmohammadi, Elena N. Kozhevnikova, Aniek van der Vaart, Ben C. Tilly, Karel Bezstarosti, Dick H. W. Dekkers, Wouter A. S. Doff, P. Jantine van de Wetering-Tieleman, Kristina Lanko, Tahsin Stefan Barakat, Tim Allertz, Jeffrey van Haren, Jeroen A. A. Demmers, Yaser Atlasi, C. Peter Verrijzer

## Abstract

Pathogenic variants in the ubiquitin-specific protease 7 (*USP7*) gene cause a neurodevelopmental disorder called Hao-Fountain syndrome. However, which of USP7’s pleiotropic functions are relevant for neurodevelopment remains unclear. Here, we present a combination of quantitative proteomics, transcriptomics and epigenomics to define the USP7 regulatory circuitry during neuronal differentiation. USP7 activity is required for the transcriptional programs that direct both differentiation of embryonic stem cells into neural stem cells, and the neuronal differentiation of SH-SY5Y neuroblastoma cells. USP7 controls the dosage of the Polycomb H2AK119ub1 ubiquitin ligase complexes ncPRC1.1 and ncPRC1.6. Loss-of-function experiments revealed that BCOR-ncPRC1.1, but not ncPRC1.6, is a key effector of USP7 during neuronal differentiation. Indeed, BCOR-ncPRC1.1 mediates a major portion of USP7-dependent gene regulation during this process. Besides providing a detailed map of the USP7 regulome during neurodifferentiation, our results suggest that USP7 and ncPRC1.1-associated neurodevelopmental disorders involve dysregulation of a shared epigenetic network.

## INTRODUCTION

Protein ubiquitylation serves a wide variety of cellular signaling functions in processes ranging from proteostasis, stress response, protein sorting to gene expression control. E.g., K48 poly- ubiquitin chains target proteins for degradation by the proteasome, while mono-ubiquitylation of histone H2A on lysine 119 (H2AK119ub1) marks Polycomb-repressed chromatin. Malfunction of the ubiquitin-proteasomal system has been implicated in various human diseases, including cancer and neurodevelopmental disorders (NDDs). Hao-Fountain syndrome (OMIM #616863) is a NDD caused by haploinsufficiency of ubiquitin-specific protease 7 (USP7).^1,2^ Affected individuals present with global developmental delay, variably impaired intellectual development with significant speech delay, behavioral anomalies, including autism spectrum disorder and attention-deficit hyperactivity disorder, eye anomalies, seizures and dysmorphic features.^1–3^ The protein deubiquitylating enzyme USP7 functions as a regulatory hub in a multi-nodal network involved in tumor suppression, the DNA damage response, Polycomb epigenetics and chromatin regulation.^4–12^ Given the pleiotropic effects of USP7 loss, it has remained unclear which of the many USP7 substrates are most relevant for neurodevelopment.

USP7 is essential for early development and viability of organisms ranging from flies to mammals.^13–14^ USP7 plays a key role in the MDM2-p53 tumor suppressor pathway, and loss of USP7 normally leads to upregulation of p53.^4,9,12^ Gene knockout (KO) studies in mice revealed that co-deletion of *Tp53* gives only a partial rescue of the *Usp7*-null phenotype.^15^ Thus, USP7 has essential biological functions during (neuro)development that are independent of p53. Indeed, USP7 substrates other than p53 play essential roles in the maintenance and expression of the genome.^7–10^ Previously, TRIM27 has been proposed as a USP7 target that is relevant for Hao-Fountain syndrome.^1^ USP7 was proposed to regulate endosomal protein recycling, through stabilization of a TRIM27-MAGEL2 ubiquitin ligase complex. However, a direct function for TRIM27 in neuronal differentiation or NDDs has not been established. In contrast, some other USP7 substrates, including several Polycomb group (PcG) proteins, have been implicated in NDDs.^16,17^ Given the central role of the Polycomb system in developmental gene control, ^18–22^ USP7 targets within this system are of particular interest for understanding its role in Hao-Fountain syndrome.

Polycomb proteins are highly conserved chromatin regulators that repress gene transcription to ensure cellular identity during development.^18–22^ Polycomb-regulated chromatin is marked by tri-methylation of H3K27 (H3K27me3) and H2AK119ub1. PcG proteins function as part of two main classes of complexes, named Polycomb repressive complex 1 (PRC1) and PRC2. The core of all mammalian PRC1 assemblies is a heterodimer of either RING1A or RING1B, paired with one of six PcG Finger (PCGF) proteins. Consequently, PRC1s are named after their incorporated PCGF, 1 through 6.^18–23^ The main activity of canonical PRC1s (cPRC1s), cPRC1.2 and cPRC1.4, is compaction of higher-order chromatin structure, while non-canonical PRC1s (ncPRC1s), ncPRC1.1, ncPRC1.3/5 and ncPRC1.6 deposit H2AK119ub1. Lastly, the key activity of PRC2 is deposition of H3K27me3 by its methyltransferase subunits EZH1 or EZH2, which is crucial for Polycomb repression. While the generality and importance of the relative order of recruitment of cPRC1, ncPRC1 and PRC2 remain controversial, there is accumulating evidence that distinct PRCs can function either independently, redundantly or cooperatively to control gene expression.^10,19,24–28^

*Drosophila* USP7 was first identified genetically as an enhancer of Polycomb repression.^13^ Subsequent studies in mammalian cells reported multiple connections between USP7 and the Polycomb system. USP7 binds, deubiquitylates and stabilizes SCML2A (a subunit of cPRC1.2/4),^29^ key subunits of ncPRC1.1 ^10,30^ and ncPRC1.6.^8,10^ Thus, both genetically and biochemically, USP7 functions as a positive regulator of the PRC1 system. Within ncPRC1.1, USP7 stabilizes BCOR, the central scaffolding- and chromatin binding subunit, KDM2B, which binds CpG islands, and the RING1A/B-PCGF1 enzymatic core.^10^ Notably, the connection between USP7 and ncPRC1.1 is conserved from flies to mammals.^23,31^ Within ncPRC1.6, USP7 stabilizes the chromatin-targeting subunits MGA and L3MBTL2, and the RING1A/B-PCGF6 enzymatic core.^8,10^ While both ncPRC1.1 and PRC1.6 play important roles in distinct tumor suppression pathways, loss-of-function of several ncPRC1.1 subunits have also been implicated in NDDs.^16,17,32,33^

It has remained unclear which of USP7’s diverse targets are most crucial during neurodevelopment. Here, we provide evidence from human cell models that USP7 function is crucial for the gene expression programs that underpin neuronal differentiation. Using a multi-omics approach, we determined the core USP7 proteomic and transcriptomic circuitry underpinning neuronal differentiation. Functional experiments established that epigenetic gene control by BCOR-ncPRC1.1 is a crucial downstream effector of USP7 during neuronal differentiation. Our results suggest that Hao-Fountain syndrome and distinct ncPRC1.1-associated NDDs involve dysregulation of a shared epigenetic network.

## RESULTS

### USP7 activity is required for neuronal differentiation

We evaluated the potential role of USP7 in neuronal differentiation in a variety of cell systems by using the potent and highly selective USP7 inhibitor FT671.^34^ We note that the addition of FT671 had no major effect on the viability of cell lines used in this study (Figure S1A). First, we tested the effect of inhibition of USP7 activity (USP7i) on the ability of H9 human embryonic stem cells (ESCs) to differentiate to neuronal stem cells (NSCs). Following transfer from expansion medium to NSC differentiation medium, H9 ESC cells readily differentiate to NSCs (Figure 1A). However, cell morphology in the presence of FT671 was strikingly different from that of control NSCs and the parental ESCs. The addition of DMSO (the solvent of FT671) alone did not affect the differentiation from ESCs to NSCs. After transfer to differentiation medium, analysis of mRNA expression showed the downregulation of pluripotency markers (*POU5f1/OCT4*, *SOX2*, *TERT*, *ZFP42/REX1*), irrespective of the presence- or absence of FT671 (Figure 1B and Data S1). However, transcriptional induction of genes associated with neural differentiation such as, *SOX3*, *PAX6*, *NES*, and *TUBΒ3*, was attenuated by FT671. Notably, expression of *PAX6*, a key regulator of neurogenesis and oculogenesis,^35^ is strongly repressed in the presence of FT671. Moreover, there was no broad induction of mesodermal or endodermal genes. Conversely, some genes that are downregulated during differentiation from ESC to NSCs, are upregulated in the presence of FT671. We conclude that USP7i blocks ESC differentiation towards NSCs (Figure 1B).

**Figure 1.**
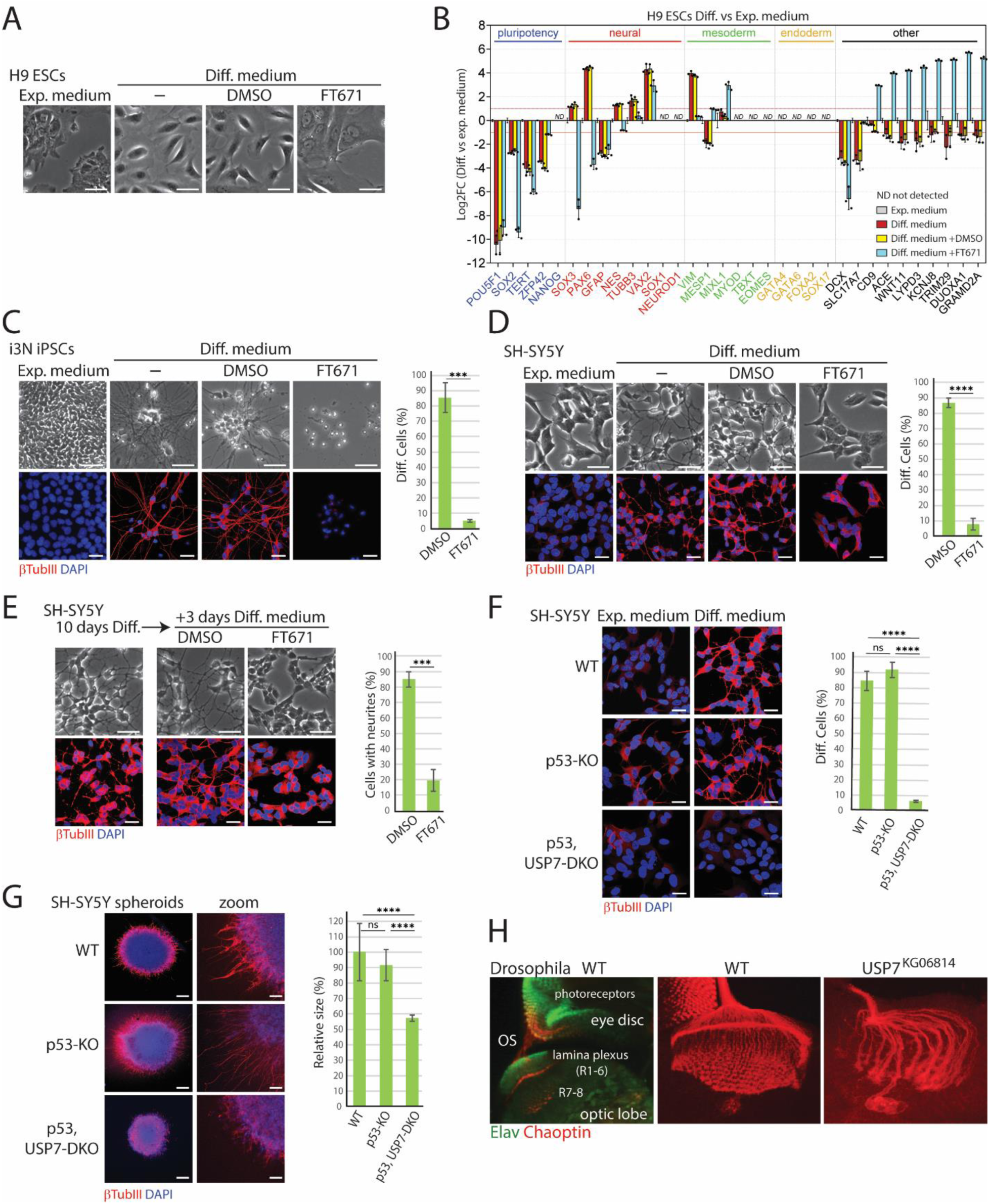
USP7 activity is crucial for neuronal differentiation of ESCs, i3N-iPSCs, SHSY-5Y neuroblastoma cells and axon development in *Drosophila* (A) Effect of USP7i on differentiation of H9 ESCs to NSCs. Phase contrast images of H9 ESCs grown in expansion medium or in NSC differentiation (Diff.) medium, either in the presence or absence of FT671. Scale bar represents 50 μm. (B) Analysis of mRNA expression of H9 ESCs grown in differentiation versus expansion (exp.) medium. Other genes refer to representative examples of the most responsive genes to USP7 inhibition during neural differentiation in H9 hESCs. Values were based on RNAseq data. Means and standard deviations (SD) were derived from three biological replicates. For RNAseq data see Data S1. (C) Effect of USP7i on differentiation of i3N-iPSCs. Phase contrast images (top panels) and IF images (bottom panels) of i3N-iPSCs grown in exp.- or differentiation medium, either in the presence or absence of FT671. The expression of βTubIII was determined by indirect IF. DNA was visualized by DAPI staining (blue). Scale bar represents 50 μm. Bar plots representing the percentage of differentiated cells, defined as having an extension from the cell body that is at least 2-times the length of the cell body. See Data S2 for quantification. Error bars represent standard deviation (SD). ***: p-value < 0.001. (D) Effect of USP7i on neuronal differentiation of SH-SY5Y cells. Analysis as described for (C). ****: p-value < 0.0001. (E) Effect of USP7i on maintenance of neuronal differentiation. SH-SY5Y cells were cultured in differentiation medium for 10 days and then shifted to media supplemented with either DMSO or FT671 for an additional 3 days. Analysis as described for (C). ***: p-value < 0.001. (F) Loss of USP7 abrogates differentiation of SH-SY5Y cells. IF images of wt, p53-KO and p53,USP7-DKO cells cultured in expansion or differentiation medium. Analysis as described for (C). See Figure S1A-F for viability assays, phase contrast images and independent clones and Data S2 for quantification. ****: p-value < 0.0001, ns: not significant. (G) Loss of USP7 compromises the formation of neurospheres. IF images of wt, p53-KO and p53,USP7-DKO cells cultured in 3D media. Scale bars represents 200 μm or 50 μm (zoom panels). See Data S2 for quantification. ****: p-value < 0.0001, ns: not significant. (H) Loss of USP7 leads to axon guidance defects in *Drosophila*. IF confocal sections of the developing visual system in wt and USP7^KG^ mutant *Drosophila* 3^rd^ instar larvae. Antibodies directed against Elav (Green) mark neurons and Chaoptin (Red) identify the photoreceptors and their axons (R1-8). OS: optic stalk.

We next tested the effect of USP7i on i3N human induced pluripotent cells (i3N-iPSCs), that expresses NEUROGENIN 2 under the control of a doxycycline-regulated promoter to drive differentiation into cortical glutamatergic i^3^Neurons.^36^ The presence of FT671 blocks neuronal differentiation of i3N-iPSCs, as judged by phase contrast microscopy and indirect immunofluorescence (IF) imaging of neuron-specific β-tubulinIII (βTubIII; Figure 1C and Data S2). Lastly, we examined the effect of USP7i on differentiation of SH-SY5Y neuroblastoma cells, which represent a convenient cell model for molecular studies of neurodifferentiation.^37^ Following treatment with retinoic acid in differentiation medium, SH-SY5Y cells stop proliferation, start to form neurites and express neuronal markers (Figure 1D and Data S2). The presence of FT671, however, strongly inhibited neuronal differentiation of SH-SY5Y cells. To test the requirement of USP7 for neuronal maintenance, we first differentiated SH-SY5Y cells, allowing formation of long neurites, followed by the addition of FT671. While βTubIII remained expressed, USP7i led to the disappearance of extended neurites (Figure 1E and Data S2). Thus, USP7 activity is required for both neuronal differentiation and neuronal maintenance. In conclusion, USP7i impedes neurodifferentiation of ESCs, iPSCs and SH-SY5Y neuroblastoma cells, indicating that deubiquitylation activity of USP7 is essential for this process.

To complement the USP7i experiments, we deleted the *USP7* gene in SH-SY5Y cells by CRISPR-Cas9 genome editing. Unfortunately, we were unable to establish stable USP7 knockout cell lines. We suspect that this failure is due to the upregulation of p53 upon loss of USP7, leading to compromised cell survival during clonal selection. Therefore, we decided to generate p53 and USP7 double knockout (DKO) SH-SY5Y cell lines (p53,USP7-DKO; Figure S1B-C). Importantly, deletion of the *TP53* gene in SH-SY5Y cells had no appreciable effect on neuronal differentiation (Figures 1F and S1D-E and Data S2). However, ablation of *USP7* in p53-KO SH- SY5Y cells blocked cell differentiation and neurite outgrowth, without affecting cell viability (Figures 1F and S1D-E). An independently generated p53,USP7-DKO clone yielded similar results (Figure S1E and Data S2). Moreover, addition of FT671 to p53-KO SH-SY5Y cells abrogated neuronal differentiation (Figure S1F and Data S2). We also found that loss of USP7 resulted in the formation of smaller neurospheres with compromised neurite outgrowth, after 3D culture of SH-SY5Y cells (Figure 1G and Data S2). We conclude that both USP7i and USP7-KO severely compromised neuronal differentiation of SH-SY5Y cells.

To determine whether the function of USP7 in neuronal differentiation is evolutionary conserved, we examined the role of USP7 in the developing *Drosophila* visual system. During 3^rd^ instar larval development, progenitor cells in the eye disk differentiate to photoreceptors that then form axons extending through the optic stalk to the optic lobe of the brain. Loss of USP7 in *Usp7^KG068^*^14^ mutant larvae^13^ abrogates the development- and guidance of axons from the photoreceptors to the optic lobe (Figures 1H and S1G). In conclusion, both inhibition and genetic ablation of USP7 demonstrate its essential role in neuronal differentiation of human (stem)cells and in developing fruit flies. We note that in the present study we focus on neuronal differentiation and did not explore the role of USP7 in alternate differentiation pathways.

### USP7 is required for transcriptional programs associated with neuronal differentiation

As a first step to uncover the molecular circuitry through which USP7 controls neurodifferentiation, we performed transcriptional profiling of H9 ESCs during differentiation towards NSCs, either in the presence or absence of FT671. Principal Component Analysis (PCA) of three independent biological replicates confirmed that the RNA-sequencing (RNA-seq) profiles of NSCs differ substantially from their parental ESCs (Figure 2A and Data S1). In agreement with the absence of NSC morphology and marker gene expression (Figures 1A-B), differentiation in the presence of FT671 leads to a transcriptional profile that differs dramatically from that of NSCs (Figure 2A). As a control, DMSO had no effect on the transcriptional profile of H9 ESC differentiation to NSCs. We identified 2,059 differentially expressed genes (DEGs) that were upregulated and 2,202 DEGs that were downregulated more than two-fold (FC>2, FDR<0.05) during differentiation from ESCs to NSCs (Figure 2B). USP7i affected ∼26% (n=526) of activated and ∼37% (n=805) of downregulated genes during differentiation (Figure 2C). Gene ontology (GO) term enrichment analysis revealed that USP7-dependent genes are mainly involved in ectoderm differentiation, axon guidance, nervous system development, muscle contraction and synaptic activity (Figure 2D). As a control, we performed transcriptional profiling of USP7i in ESC cells cultured in expansion medium. We observed only ∼10% overlap of DEGs in response to USP7i in ESCs with USP7-dependent DEGs following differentiation to NSCs (Figure S2A-B and Data S1). Indeed, GO term enrichment analysis of DEGs after USP7i in ESCs reflected very distinct biological pathways (e.g. cholesterol metabolism or herpes simplex virus infection) from USP7i in differentiation medium (Figure S2C-D).

**Figure 2.**
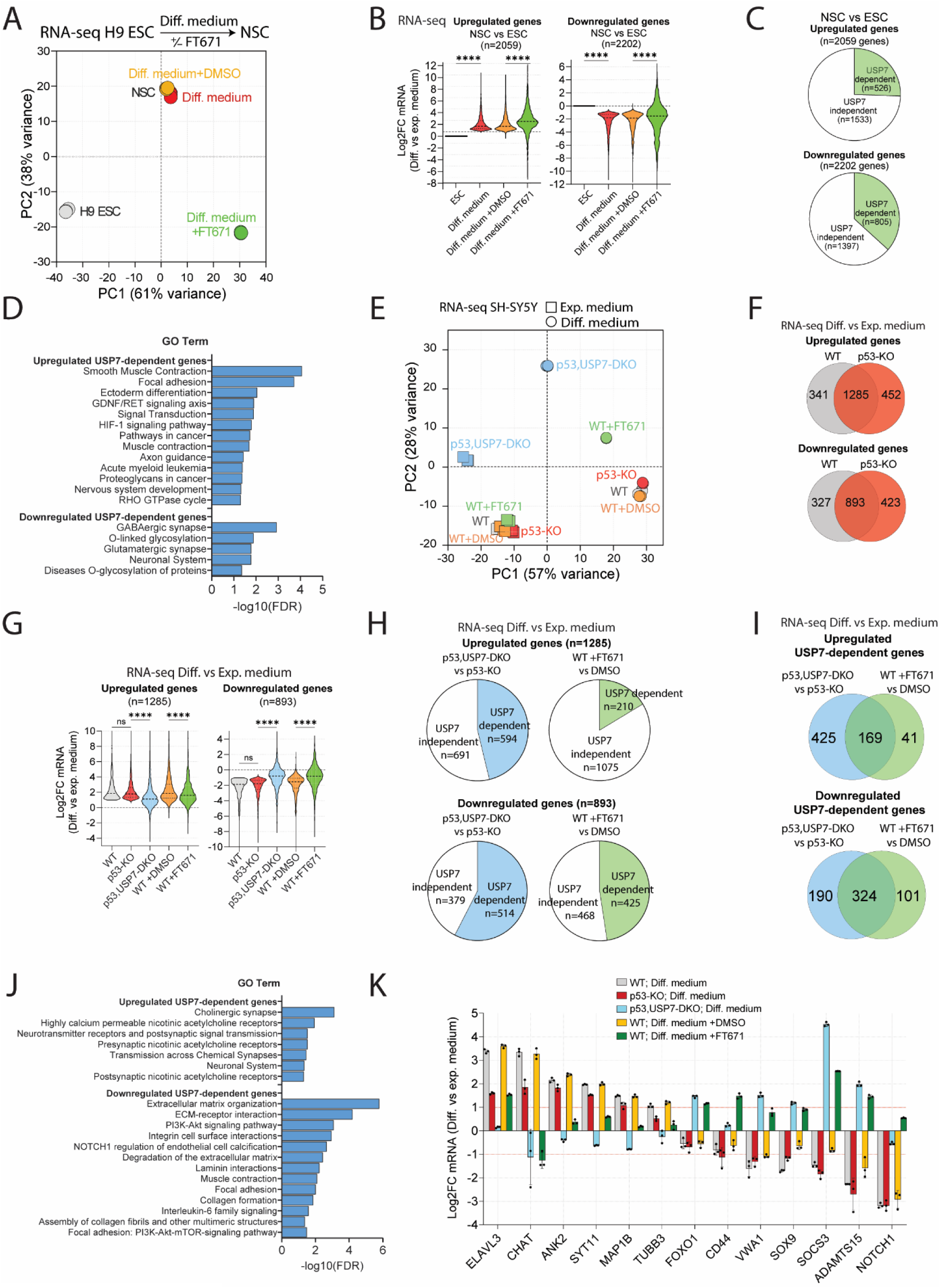
**USP7 controls the transcriptional program associated with neural differentiation.** (A) PCA of RNA-seq data from H9 ESCs grown in expansion medium or differentiation medium the absence or presence of DMSO or FT671. For each condition 3 independent biological replicates were used. For RNAseq data see Data S1. (B) Violin plots of DEGs in H9 ESCs vs differentiated NSCs. Expressions of these genes are also shown in H9 ESCs cultured in NSC differentiation medium supplemented with DMSO or FT671. Means were derived from 3 biological replicates per condition. ****: p-value < 0.0001. (C) Pie chart showing the percentage of DEGs that are significantly affected by FT671 treatment (USP7-dependent). (D) GO Term analysis (KEGG) of USP7-dependent genes identified in (C). (E) PCA of RNA-seq data from wt (+/- DMSO or FT671), p53-KO and p53,USP7-DKO SH- SY5Y cells before and after 10 days of culture in differentiation medium (3 biological replicates per condition). For RNAseq data see Data S3. (F) Venn diagrams showing the overlap of DEGs in wt or p53-KO cells before and after 10 days of neural differentiation. The intersect represent p53-independent DEGs. (G) Violin plots of p53-independent DEGs during neuronal differentiation identified in Figure 2F. Fold change reflects gene expression in differentiation medium vs expansion. Means were derived from 3 biological replicates per condition. ****: p-value < 0.0001. (H) Pie chart showing the percentage of p53-independent neuro-differentiation DEGs (Figure 2F) that are significantly affected by loss of USP7 (p53,USP7-DKO vs p53-KO) or USP7i (FT671 vs DMSO). (I) Venn diagrams showing the overlap of neuro-differentiation DEGs affected by *USP7* KO and USP7 inhibition. (J) GO term analysis (KEGG) of high confidence USP7-dependent genes identified in Figure 2I. (K) Barplot showing examples of USP7-dependent neural differentiation genes. Means and SDs were derived from 3 biological replicates. For RNAseq data see Data S3.

Next, we extended our transcriptional profiling to differentiating SH-SY5Y cells. We compared the RNA-seq profiles of wt, p53-KO and p53,USP7-DKO cells before and after neuronal differentiation. Additionally, we analyzed the transcriptional impact of USP7i on wt SH-SY5Y cells during neuronal differentiation. Thus, we could assess both p53-dependent and p53- independent functions of USP7. PCA of sequenced mRNA, isolated from three independent biological replicates, revealed that wt and p53-KO cells exhibited largely similar behavior during neuronal differentiation, sharing 1,285 upregulated and 893 downregulated genes (Figures 2E-G and Data S3). In contrast, the transcriptional differentiation program in p53,USP7-DKO cells was strongly impaired. Likewise, USP7i caused the transcription profile of wt SH-SY5Y cells grown in differentiation medium to shift away from differentiated control cells, towards that of p53,USP7-DKO cells (Figure 2E). Taken together, these results show that USP7 plays a major role, independent of p53, in the regulation of the transcriptional program driving SH-SY5Y cell neuronal differentiation.

To define the high-confidence USP7-dependent core transcriptional circuitry during neuronal differentiation, we identified genes whose expression was affected both by USP7i (in wt SH-SY5Y cells) and by deletion of *USP7* (in p53-KO cells). This analysis yielded 169 upregulated and 324 downregulated USP7-dependent genes during neuronal differentiation (Figures 2H and I). USP7-dependent genes that are induced during neuronal differentiation are predominantly associated with GO terms related to neurogenesis, and include *ELAVL3*, *CHAT*, *ANK2*, *SYT11*, *MAP1B* and *TUBB3* (Figures 2J-K). Genes downregulated during neuronal differentiation, but aberrantly expressed due to loss of USP7 function, were enriched for GO terms associated with extracellular matrix organization, PI3K/AKT, FOXO1 and NOTCH signaling pathways. Notably, FOXO1 has been implicated in the promotion of neural cell death.^38^ Moreover, SOX9, a key transcriptional regulator of morphogenesis and a suppressor of neurogenesis, is repressed during neuronal differentiation in a USP7-dependent manner. Taken together, our analysis identifies the USP7-dependent transcriptional circuitries of ESC differentiation to NSCs, and of neuronal differentiation of SH-SY5Y cells.

### USP7 controls the dosage of ncPRC1.1, ncPRC1.6 and TRIM27 during neuronal differentiation

Because USP7 is a deubiquitylating enzyme that can regulate protein levels at the post-translational level, we also monitored USP7-dependent changes in the SH-SY5Y cell proteome during neuronal differentiation. To determine relative protein abundances, we used data-independent acquisition mass spectrometry (DIA-MS) combined with label-free quantification (LFQ). First, we determined the effect of USP7i on the proteome of SH-SY5Y cells grown for 10 days in either expansion- or differentiation medium. PCA on 4,914 protein entries that had valid intensity values under all conditions and for all three independent biological replicates, revealed a clear effect of FT671 on the proteome, especially under differentiation condition (Figure 3A and Data S4). Selection of proteins with similar dynamic abundance profiles allowed the identification of those that are up- or downregulated during neuronal differentiation in a USP7-dependent manner (Figures 3B-D and S3). GO-term enrichment analysis of the top 100 USP7-dependent upregulated proteins revealed that they are overwhelmingly involved in neuronal development (Figures 3B-C and S3). In contrast, the top 100 USP7-dependent down-regulated proteins are involved in extracellular vesicle function and metabolic processes (Figures 3D-E and S3). Additionally, we compared the proteomic profiles of wt, p53-KO and p53,USP7-DKO SH-SY5Y cells before and after neural differentiation (Figures 3F-J and S3 and DataS5). PCA on 5,657 protein entries that had valid intensity values in all replicates under all conditions, showed that the proteomes of p53-KO cells cluster close to those of wt cells (Figure 3F). In contrast, the proteomes of p53,USP7-DKO cells differed substantially from wt and p53-KO cells, both in expansion and in differentiation medium. Similarly to the USP7i experiments, GO analysis identified USP7- dependent induction of proteins involved in neuronal development and axon formation, while extracellular vesicle and metabolic pathways were downregulated in a USP7-dependent manner (Figures 3G-J and S3). Collectively, the results from cellular differentiation, transcriptomic and proteomic profiling thus establish an essential function for USP7 in neuronal differentiation and provide detailed information on the molecular circuitry involved.

**Figure 3.**
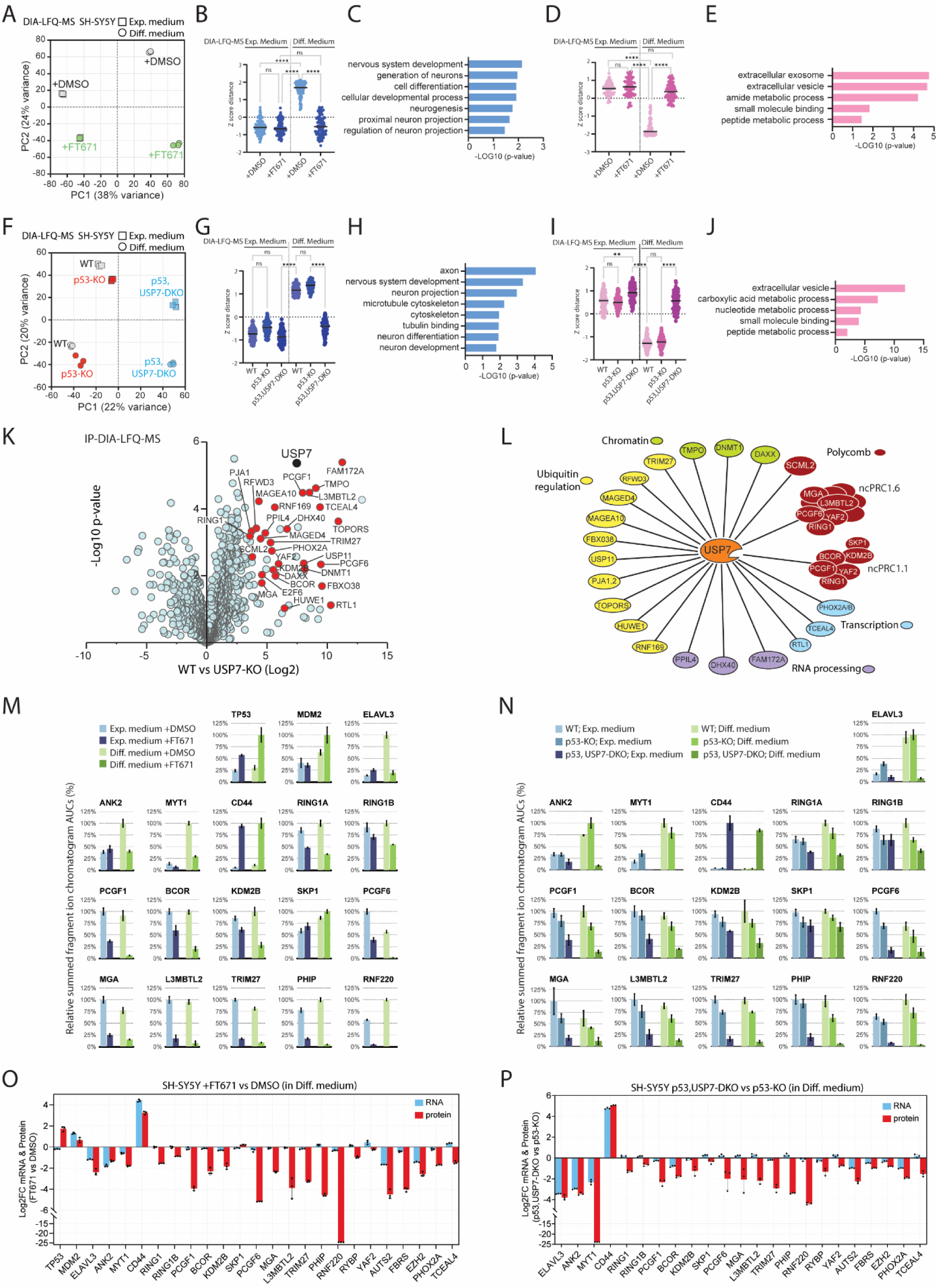
**Quantitative analysis of the USP7 protein network during neuronal differentiation.** (A) Impact of USP7i on the SH-SY5Y cell proteome during neural differentiation. PCA of DIA-MS data from SHSY-5Y cells grown for 10 days in expansion or in differentiation medium in the presence or absence of FT671 (3 biological replicates per condition). PCA was on 4,914 protein entries that had valid intensity values under all conditions and for all 3 independent biological replicates (out of 8,568 protein entries in total). For the complete set of protein profiles see Figure S3. For the complete data set see Data S4. (B) Plot of top 100 proteins that are normally induced during differentiation but fail to do so in the presence of FT671. ****: p-value < 0.0001, ns: not significant. (C) GO analysis of USP7-dependent proteins induced during differentiation selected in (B). (D) Plot of top 100 proteins that are normally repressed during differentiation but fail to do so in the presence of FT671. ****: p-value < 0.0001, ns: not significant. (E) GO analysis of USP7-dependent proteins repressed during differentiation selected in (D). (F) Impact of loss of USP7 on the SHSY-5Y cell proteome during neural differentiation. PCA of DIA-MS data from wt, p53-KO and p53,USP7-DKO SH-SY5Y cells grown for 10 days in expansion- or in differentiation medium (3 biological replicates per condition). PCA was on 5,657 protein entries that had valid intensity values under all conditions and for all 3 independent biological replicates (out of 8,241 protein entries in total). For the complete set of protein profiles see Figure S3. For the complete data set see Data S5. (G) Plot of top 100 USP7-dependent proteins that are normally induced during differentiation. ****: p-value < 0.0001, ns: not significant. (H) GO analysis of USP7-dependent proteins induced during differentiation selected in (G). (I) Plot of top 100 USP7-dependent proteins that are normally repressed during differentiation but fail to do so in the absence of USP7. ****: p-value < 0.0001, **: p-value < 0.01, ns: not significant. (J) GO analysis of USP7-dependent proteins repressed during differentiation selected in (I). (K) Proteins associated with endogenous USP7 purified from SH-SY5Y cells. Volcano plot in which the permutation-based false discovery rate corrected t-test difference of the LOG2 iBAQ values of endogenous USP7 IPs from wt SH-SY5Y cells versus control IPs from pP53,USP7-DKO cells are plotted against the negative LOG10 p-value of the test. The analysis is based on three independent biological replicate experiments. High-confidence USP7-interacting proteins are shown in red. Grey circles depict common contaminants like ribosomal proteins or insignificant hits. For the complete data set see Data S6. (L) A schematic summary of high confidence interactions. PRC1 subunits are depicted as part of established complexes. (M) PRM analysis of relative protein abundance in wt SH-SY5Y cells grown for 10 days in expansion- or in differentiation medium in the presence or absence of FT671 (3 biological replicates per condition). Chromatographic areas under the curve (AUCs) of peptides were summed to yield their respective protein quantities, which were subsequently averaged over three replicates and are shown as a percentage (± SD) of the largest resulting mean. For additional proteins see Figure S4A. For the complete data set see Data S7. (N) PRM analysis of relative protein abundance in wt, p53-KO and p53,USP7-DKO SH-SY5Y cells grown for 10 days in expansion- or in differentiation medium (3 biological replicates per condition). For additional proteins see Figure S4B. For the complete data set see Data S7. (O) Barplot of changes in mRNA- (blue) and protein (red) levels in wt SH-SY5Ycells grown in differentiation medium in the presence or absence of FT671. Values based on RNA-seq- and PRM data. (P) Barplot of changes in mRNA- (blue) and protein (red) levels in wt, p53-KO or p53,USP7-DKO SH-SY5Y cells grown in differentiation medium. Values based on RNA-seq- (Data S3) and PRM data (Data S7). Means and SDs were derived from 3 biological replicates.

To determine its interacting partners in SH-SY5Y cells, we immune-purified (IPed) endogenous USP7 from whole cell extracts followed by mass spectrometry. As a negative control, we used anti-USP7 antibodies for IPs from p53,USP7-KO SH-SY5Y cells. IPs were performed on three biological replicates, followed by on-bead digestion and protein identification by mass spectrometry. To identify significant interacting proteins, we determined the permutation-based false discovery rate corrected t-test of protein enrichment in IPs from wt versus USP7-KO cells (Figure 3K and Data S6). This analysis confirmed previously identified high-confidence USP7- interacting proteins including BCOR, DAXX, DNMT1, PCGF1, PCGF6, TCEAL4 and TRIM27, but also provided support for novel and less-established partners, such as e.g., FAM172A, TOPORS, PHIP, PPIL4, PJA1, RNF169, RNF220, TMPO and PHOX2A (see also references (5–11)). The USP7 interactome largely comprises proteins involved in nuclear ubiquitin regulation, chromatin, transcription and RNA processing (Figure 3L). SCML2, ncPRC1.1 and ncPRC1.6 are prominent USP7-interacting components of the Polycomb system, whereas we did not detect association of other PcG proteins.

Next, we used parallel reaction monitoring (PRM)-MS, as a comprehensive and quantitative method to determine the impact of USP7 on the abundance of selected markers of neuronal differentiation and interacting proteins (Figures 3M-N and S4 and Data S7). We found that USP7i stabilized p53 but did not lead to decreased levels of MDM2 in wt SH-SY5Y cells (Figure 3M). In both wt and p53-KO cells, growth in differentiation media resulted in a strong induction of neuronal proteins such as ELAVL3, ANK2 and MYT1 (Figures 3M-N). Consistent with the crucial role of USP7 in neuronal differentiation, this induction was blocked by USP7i and by USP7-KO. CD44 is an example of a protein that is strongly upregulated upon loss of USP7 activity, independent of neuronal differentiation. USP7 is required for the stabilization of key ncPRC1.1 subunits PCGF1, BCOR and KDM2B. In contrast, SKP1, which is part of a multitude of other protein assemblies, is not affected by loss of USP7 activity. Through stabilization of PCGF6, MGA and L3MBTL2, USP7 also controls the functional integrity of ncPRC1.6. Additionally, we observed a strong dependency on USP7 for other high confidence substrates such as TRIM27, PHIP and RNF220 (see Figure S4 for additional USP7 targets). We note that the stabilizing effect of USP7 on its substrates is generally independent of neuronal differentiation. Moreover, in addition to regulating protein levels, deubiquitylation by USP7 may also modulate protein function without affecting its abundance.

To determine the relative importance of gene transcription versus post-translational regulation by USP7, we compared changes in mRNA and protein levels in cells grown in differentiation medium (Figures 3O-P). As expected, upregulation of p53 upon inhibition of USP7 occurs at the post-translational level. The increased amount of p53 explains the mild activation of *MDM2* transcription (Figure 3O). In agreement with previous results,^11^ we do not observe a downregulation of MDM2 protein due to USP7i. The aborted induction of neuronal markers such as ELAVL3 and ANK2 in the absence of USP7 activity, appears to be primarily due to reduced mRNA levels. However, USP7-dependent stabilization of RING1A and RING1B, PCGF1, BCOR, KDM2B, PCGF6, MGA and L3MBTL2 was not accompanied by appreciable changes in mRNA levels. Thus, ncPRC1.1 and ncPRC1.6 are likely direct targets of USP7 whose levels are regulated at the post-translational level. Likewise, TRIM27, PHIP, RNF220, TCEAL4 and some additional USP7-binding proteins are stabilized by USP7 at the post-translational level. For some factors, such as MYT1, PHOX2A, AUTS2 and FBRS, we observed both transcriptional and post-translational changes. In the absence of observable binding to USP7, it remains undetermined how these proteins might be regulated. Taken together, these results provide a detailed description of the USP7-dependent transcriptomic and proteomic changes that underpin neuronal differentiation. We established that USP7 directly binds and stabilizes ncPRC1.1, ncPRC1.6, TRIM27 as well as other substrates.

### Heterozygous loss of USP7 impedes the neuronal differentiation program

Above, we used inhibition of USP7 activity by FT671 or full USP7 KOs to identify its substrates and downstream transcriptional network during neuronal cell differentiation. Because Hao-Fountain syndrome is caused by haploinsufficiency of USP7, we next generated a USP7 heterozygous (USP7^het^) clone that expresses USP7 at approximately half its level in wt SH-SY5Y cells (Figures 4A-B). Strikingly, this ∼50% reduction in USP7 levels was mirrored by a ∼50% decrease in neurite formation (Figure 4C and Data S2). Thus, the efficiency of neuronal differentiation of SH-SY5Y cells is highly dependent on USP7 dosage. To determine the impact of USP7 heterozygosity on gene expression, we compared the transcriptional profiles of USP7^het^ and wt cells. PCA revealed notable differences between the transcriptomes of USP7^het^ and WT cells cultured in either expansion- or differentiation medium (Figure 4D and Data S8). We next focused on USP7-dependent genes associated with neuronal differentiation that we identified earlier in USP7i and USP7-KO cells (Figure 2). Analysis of these genes in USP7^het^ cells showed intermediate, yet significant, changes in gene expression when compared to the effects of USP7i or full USP7 deletion (Figures 4E). While not as strong as in USP7-KO or USP7i cells, the expression of key developmental genes during neuronal differentiation is clearly affected in USP7^het^ cells (Figures 4E).

**Figure 4.**
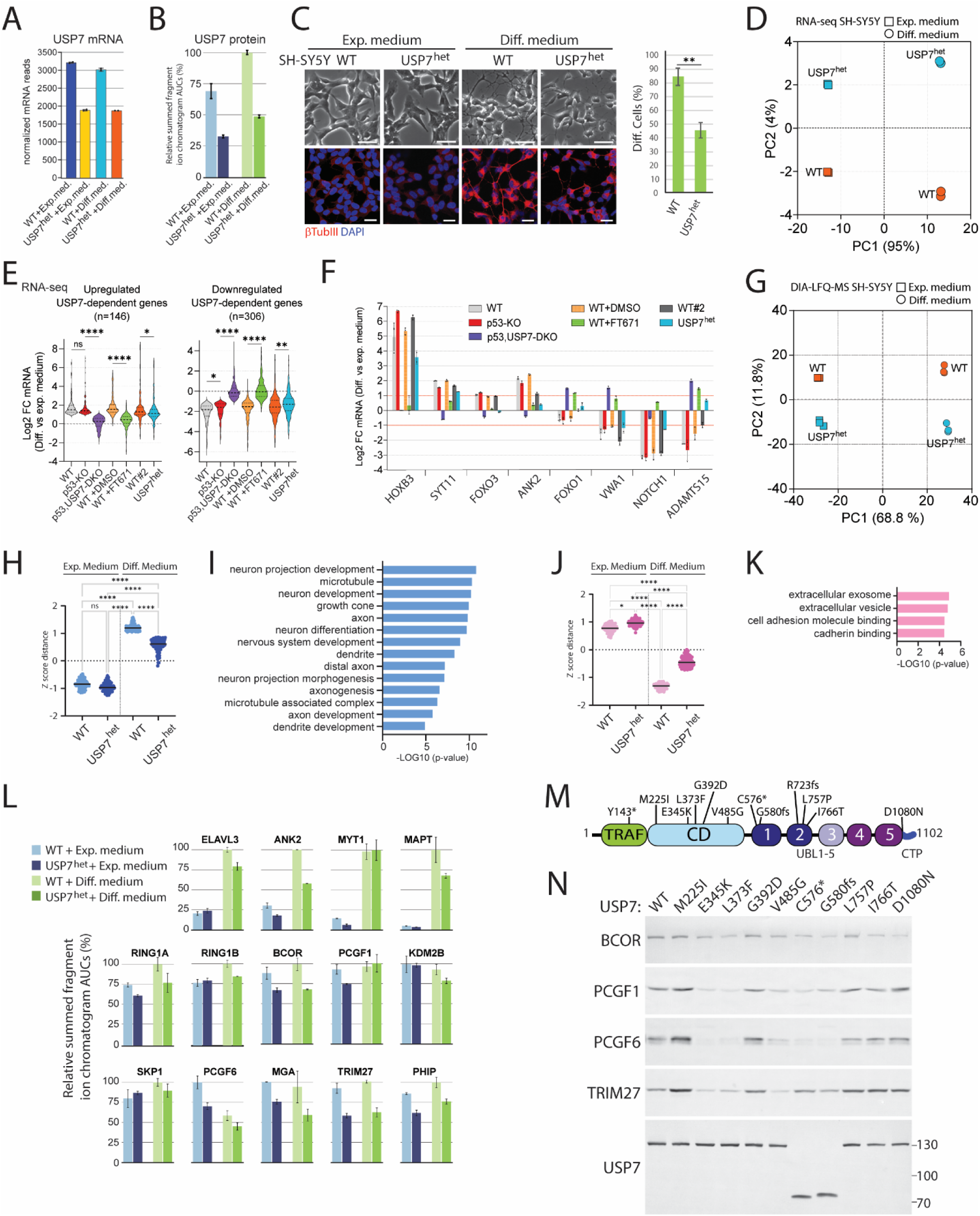
**Heterozygous loss of USP7 impedes the neuronal differentiation program of SH- SY5Y cells.** (A) USP7 mRNA expression in wt and USP7^het^ SH-SY5Y cells cultured in expansion or differentiation medium. Values were based on RNAseq data. Means and SDs were derived from 3 biological replicates. For RNAseq data see Data S8. (B) Targeted detection by PRM-MS of USP7 protein in wt and USP7^het^ SH-SY5Y cells cultured in expansion or differentiation medium. Values were based on summed chromatographic areas under the curve (AUCs) of unique tryptic peptides of USP7. Means and SDs were derived from 3 biological replicates. (C) Analysis of neuronal differentiation of wt and USP7^het^ SH-SY5Y cells, as described for Figure 1C. See Data S2 for quantification. **: p-value < 0.001. (D) PCA of RNA-seq data from wt and USP7^het^ SH-SY5Y cells before and after 10 days of culture in differentiation medium (2 biological replicates per condition). For RNAseq data see Data S8. (E) Violin plots of DEGs during neuronal differentiation identified in Figure 2I. Fold change reflects gene expression in differentiation- versus expansion medium. Means were derived from 3 biological replicates per condition. ****: p-value < 0.0001; ** p-value < 0.001; *: p-value < 0.05. wt#2 refers to RNA samples from wt cells isolated in parallel with RNA from USP7^het^ cells. (F) Barplot showing examples of USP7-dependent DEGs associated with neural differentiation. Means and SDs were derived from RNA-seq data using 2 (wt & USP7^het^) or 3 biological replicates. (G) Impact of USP7 heterozygosity on the SH-SY5Y cell proteome during neural differentiation. PCA of DIA-MS data from wt and USP7^het^ cells grown for 10 days in either expansion or in differentiation medium (3 biological replicates per condition). PCA was on 4,730 protein entries that had valid intensity values under all conditions and for all 3 independent biological replicates (out of 6,548 protein entries in total). For the complete data set see Data S9. (H) Plot of top 100 proteins that are induced during differentiation of wt cells but display attenuated induction in USP7^het^ cells. ****: p-value < 0.0001, ns: not significant. (I) GO analysis of USP7-dependent proteins induced during differentiation, selected in (H). (J) Plot of top 100 proteins that are repressed during differentiation of wt cells, but that are less repressed in USP7^het^ cells. ****: p-value < 0.0001, *: p-value < 0.05, ns: not significant. (K) GO analysis of USP7-dependent proteins repressed during differentiation, selected in (J). (L) PRM-MS analysis of relative protein abundance in wt and USP7^het^ SH-SY5Y cells grown for 10 days in expansion or in differentiation medium (3 biological replicates per condition). Chromatographic areas under the curve (AUCs) of peptides were summed to yield their respective protein quantities, which were subsequently averaged over three replicates and are shown as a percentage (± SD) of the largest resulting mean. For additional proteins see Figure S5A. For the complete data set see Data S10. (M) Hao-Fountain syndrome associated variants (from Fountain et al., 2019). Functional domains of USP7 are indicated: TRAF domain, CD: catalytic domain, UBL1-5, CTP: C-terminal peptide. (N) The impact of Hao-Fountain syndrome-associated patient mutations on USP7 function. Immunoblot analysis of USP7-KO HEK293T cells co-transfected with vectors expressing either Flag-tagged wt or the indicated USP7 mutants, in combination with vectors expressing either BCOR, PCGF1, PCGF6 or TRIM27. See also Figures S5B-D.

PCA on 4,730 protein entries, identified by DIA-MS, that had valid intensity values under all conditions and for all 3 independent biological replicates (out of 6,548 identified protein entries in total) revealed clear differences between the proteomes of USP7^het^ and wt cells (Figure 4G and Data S9). GO-term analysis of the top 100 proteins that are induced during differentiation of wt cells, but display attenuated upregulation in USP7^het^ cells, revealed that they are predominantly involved in neuritogenesis and neuronal differentiation (Figures 4H-I). USP7-dependent proteins that are down-regulated during growth in differentiation medium are mainly involved in extracellular processes and cell adhesion (Figures 4J-K). Thus, although the effects are milder, USP7 heterozygosity significantly affects similar neurodifferentiation pathways as complete ablation of USP7 or USP7i. Next, we took advantage of the ability of PRM-MS to measure small changes in relative protein abundances, to determine the effect of USP7 heterozygosity on a selection of USP7 substrates and neuronal differentiation proteins. PRM-MS revealed that different targets displayed different sensitivities to the reduced level of USP7 in USP7^het^ cells (Figures 4L and S5A and Data S10). Notably, the levels of substrates like BCOR, MGA, PCGF6 and TRIM27 were reduced in USP7^het^ compared to wt cells. In conclusion, a ∼50% reduction in the level of USP7 causes small changes in the levels of many of its substrates, which collectively have a substantial impact on gene transcription and neuronal differentiation.

Lastly, we tested the impact on USP7 function of a series of 12 missense and truncating USP7 variants (Figure 4M), which were identified in patients diagnosed with Hao-Fountain syndrome.^2^ First, we determined the protein levels of Flag-tagged USP7 variants following their expression in HEK293T USP7-KO cells (Figure S5B). The expression of USP7-Y143* and USP7- R723fs was severely compromised, which might be due to nonsense mediated decay or protein instability, and these variants were therefore not analyzed further. USP7-C576*, USP7-G580fs and USP7-I766T were expressed at reduced levels, while the protein levels of the remaining variants were comparable to that of wt USP7. Next, we verified the subcellular localization of variant USP7. In contrast to endogenous USP7, which is predominantly nuclear (Figure S5C), truncation mutants USP7-C576* and G580fs were mainly localized in the cytoplasm (Figure S5D). Thus, some USP7 variants that are associated with Hao-Fountain syndrome affect protein levels or subcellular localization. To determine the effect of mutations on USP7 activity, we co-transfected vectors expressing either wt USP7 or the indicated variants, with vectors expressing selective substrates into HEK293T USP7-KO cells. We adjusted the amounts of USP7-expressing vectors to yield comparable levels of USP7 expression (Figure 4N). The ability to stabilize BCOR was compromised in 8 USP7 variants, whereas the activity of USP7-M225I and G392D was comparable to that of wt USP7 (Figure 4N). Stabilization of PCGF1 and PCGF6 was compromised in USP7-E345K, L373F, V485G, C576* and G580fs, but not in the remaining variants. Levels of TRIM27 were notably lower when co-expressed with USP7-E345K, L373F and V485G, but the impact of C576* and G580fs was milder than observed for PCGF1 and PCGF6. Notably, USP7- M225I and USP7-G392D appear to be more active than wt USP7 in the stabilization of some substrates. Thus, they might represent gain-of-function variants.

In summary, these results establish that neuronal differentiation is highly dependent on USP7 dosage. Heterozygosity of USP7 causes modest but significant changes in gene expression that result in a ∼50% impairment of neuronal differentiation. The stabilization of USP7 substrates is affected by variants associated with Hao-Fountain syndrome. However, different substrates display distinct sensitivities to some of the USP7 variants. Although a more rigorous analysis is required, some missense mutations associated with Hao-Fountain syndrome, such as USP7-M225I and USP7-G392D, might create hyperactive gain-of-function variants of USP7.

### BCOR-ncPRC1.1, but not ncPRC1.6 or TRIM27, is required for neuronal differentiation

Next, we set out to determine which of the USP7 targets are crucial for neuronal differentiation. Because of its crucial role in developmental gene control, we first tested the requirement of the Polycomb system for neuronal differentiation of SH-SY5Y cells. Deletion of either RING1A or RING1B caused an approximately 50% reduction in neurite formation of SH-SY5Y cells grown in differentiation medium (Figures 5A and S6 and Data S2). Thus, neuronal differentiation of SH- SY5Y cells is highly sensitive to reduced dosage of PRC1. In spite of screening over 200 clones, we failed to obtain lines that were deficient for both RING1A and RING1B, indicating that loss of both affects SH-SY5Y cell viability. Next, we tested the relative importance of USP7 targets ncPRC1.1 and ncPRC1.6 in neuronal differentiation. Loss of BCOR, the central architectural subunit of ncPRC1.1, abrogated cell differentiation and neurite outgrowth (Figures 5A and S6 and Data S2). In contrast, loss of PCGF6 had no impact on neuronal differentiation. Thus, while ncPRC1.1 and ncPRC1.6 both depend on USP7 for their functional integrity, only ncPRC1.1 is required for neuronal differentiation of SH-SY5Y cells. While TRIM27 has been linked to Hao-Fountain syndrome,^1^, its deletion had no appreciable effect on neurodifferentiation of SH-SY5Y cells (Figures 5A and S6 and Data S2). Loss of another USP7 target protein, PHIP/BRWD2,^8^ substantially reduced neuritogenesis. We conclude that BCOR-ncPRC1.1 is a crucial USP7 target during neuronal differentiation of SH-SY5Y cells, although other substrates, such as PHIP, are also involved in this process.

**Figure 5.**
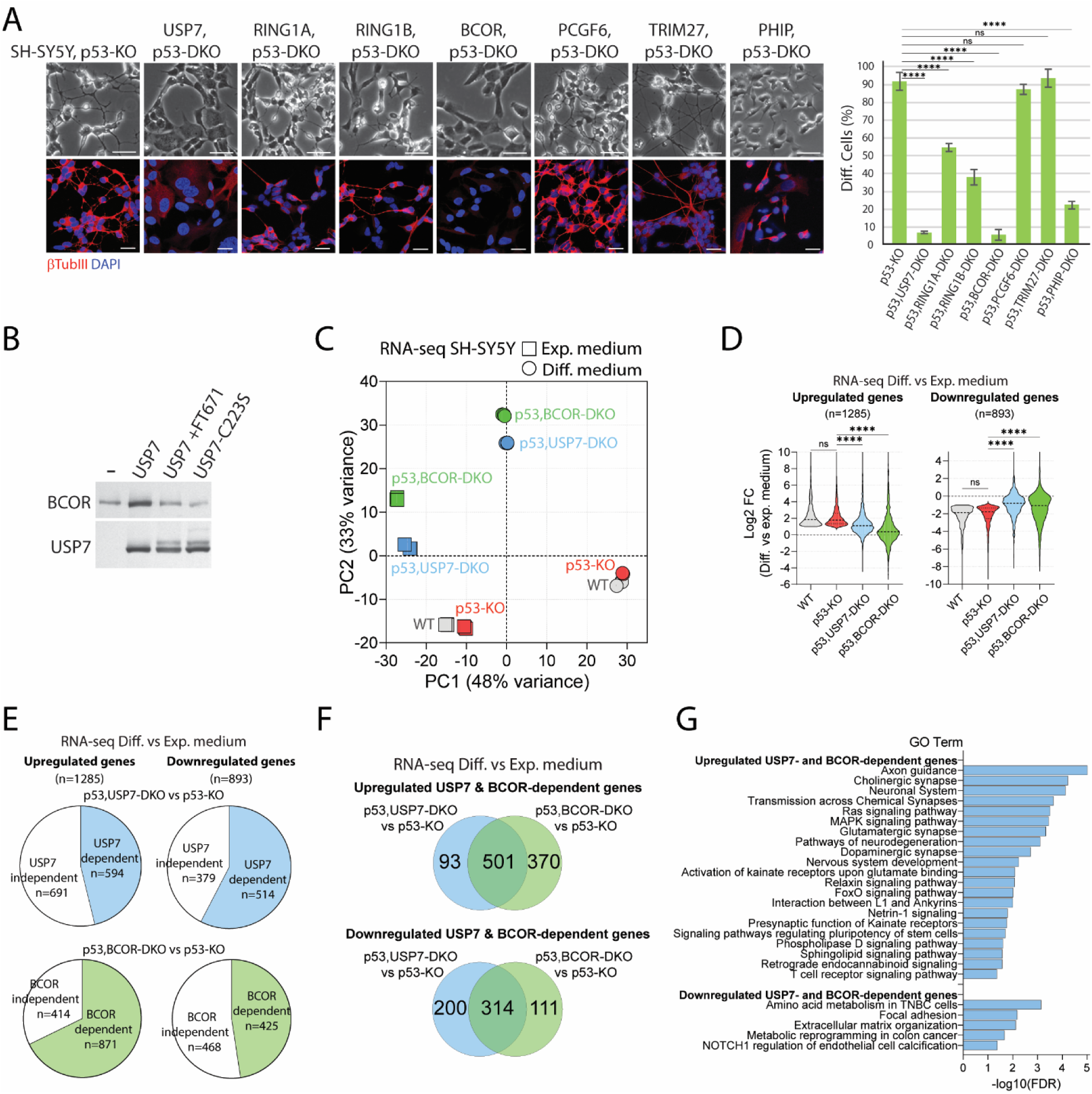
BCOR is a major downstream effector of USP7 function during neuronal differentiation. (A) Effect of deletion of several USP7 targets on neuronal differentiation of p53-KO SH-SY5Y cells. Analysis as described for Figure 1C. Scale bar represents 50 μm. For additional images and the analysis of independent clones see Figure S6A-B. ****: p-value < 0.0001, ns: not significant. See Data S2 for quantification. (B) USP7 activity is required for the stabilization of BCOR. Immunoblot analysis of HEK293T USP7-KO cells co-transfected with a vector expressing GFP-BCOR, in combination with either an empty vector (-), a vector expressing wt USP7 (in the presence or absence of FT671) or a vector expressing USP7-C223S. (C) PCA of RNA-seq data from wt p53-KO, p53,USP7-DKO and p53,BCOR-DKO SH-SY5Y cells before and after 10 days of culture in differentiation medium (3 biological replicates per condition). For RNA-seq data see Data S3. (D) Violin plots of DEGs during neuronal differentiation identified in Figure 2F. Fold change reflects gene expression in differentiation- versus expansion medium. Means were derived from 3 biological replicates per condition. ****: p-value < 0.0001; ns: not significant. (E) Pie chart showing the percentage of DEGs associated with neuro-differentiation (Figure 2F) that are significantly affected by the loss of USP7 (p53,USP7-DKO vs p53-KO) or BCOR (p53,BCOR-DKO vs p53-KO). (F) Venn diagrams showing the overlap of neuro-differentiation DEGs affected by loss of USP7 or BCOR, identified in (C). (G) GO term analysis (KEGG) of high confidence USP7- and BCOR-dependent genes identified in (E).

### BCOR is a major downstream effector of USP7 function during neuronal differentiation

The results from USP7i experiments (Figure 3M-O) suggested that substrate stabilization by USP7 is dependent on its deubiquitylation activity. As an additional test for BCOR, we co-transfected a vector expressing BCOR in HEK293T USP7-KO cells with vectors expressing either no protein (-), wt USP7 or catalytically inactive USP7-C223S (Figure 5B). USP7, but not USP7-C223S stabilized BCOR. Moreover, the addition of FT671 inhibited the stabilization of BCOR by wt USP7. Collectively, our results show that stabilization of BCOR by USP7 is dependent on its enzymatic activity.

To define the role of BCOR as a downstream effector of USP7 function, we compared the transcriptional profiles of p53,BCOR-DKO, p53,USP7-DKO and p53-KO SH-SY5Y cells during neuronal differentiation. Remarkably, PCA of the RNA-seq profiles showed that the transcriptional impact of loss of BCOR largely phenocopied that of USP7 deletion (Figure 5C). Loss of BCOR affects 68% of genes (871 out of 1,285 induced DEGs) that are activated during neuronal differentiation in both WT and p53-KO cells (as defined in Figure 2F), and 48% of genes (425 out of 893 repressed DEGs) that are repressed during this process (Figure 5D-E). Thus, BCOR has a major impact on the gene regulatory program of neurodifferentiation, phenocopying loss of USP7 function. A comparison of USP7- and BCOR-dependent DEGs (FC>2, FDR< 0.05), revealed that the expression of 73% of genes affected by loss of USP7 are also impaired by deletion of BCOR (815 out of 1,108 DEGs; Figure 5F). These USP7- and BCOR-dependent genes are predominantly enriched for GO-terms associated with neural development, such as axon guidance and synapse function (Figure 5G). Collectively, results from cell differentiation experiments, transcriptomic and proteomic analysis strongly support the notion that BCOR-ncPRC1.1 is a major downstream effector of USP7 function during neuronal differentiation.

### BCOR binds key neuronal differentiation genes

To establish which genes are targeted directly by BCOR in a USP7-dependent manner, we determined its genomic distribution in wt SH-SY5Y cells grown in differentiation medium for 3 days, either in the presence or absence of FT671. In parallel, we determined the effect of USP7i on the Polycomb chromatin marks H2AK119ub1 and H3K27me3. Chromatin immunoprecipitation with reference exogenous genome (ChIP-Rx) using *Drosophila* chromatin as a spiked-in normalization reference, allowed a direct and quantitative comparison between histone modifications in the presence or absence of USP7i. Our ChIP-Rx-seq analysis identified 6,247 BCOR, 23,788 H2AK119ub1 and 17,329 H3K27me3 high-confidence peaks in DMSO-treated control cells (Figure 6A). Analysis of the distribution of H2AK119ub1 and H3K27me3 revealed three classes of genomic loci (Figure 6B): (1) 15,384 loci marked by H2AK119ub1 but devoid of H3K27me3, (2) 12,118 loci only marked by H3K27me3 and (3) 8,767 loci marked by both H2AK119ub1 and H3K27me3. Thus, while H2AK119ub1 and H3K27me3 co-exist at about one third of their loci, they are not globally coincident. BCOR binding sites overlap overwhelmingly with H2AK119ub1 marked regions, but not with H3K27me3 in the absence of H2AK119ub1 (Figures 6A-B). Notably, BCOR peaks were predominantly enriched at CpG islands, promoters and 5’UTRs (Figure 6C). In line with its destabilizing effect on BCOR- ncPRC1.1 (Figures 3M-N), the presence of FT671 eliminated BCOR binding to chromatin (Figures 6B and D). However, the effect of USP7i on chromatin binding by BCOR appears to be more dramatic than its effect on BCOR protein levels. USP7i also caused a notable drop in the levels of genomic H2AK119ub1, with minor effects on global H3K27me3. Note that the global reduction in H2AK119ub1 is the result of reduced levels of both BCOR-ncPRC1 and ncPRC1.6 (Figures 3M-N), as well as potential indirect mechanisms. The impact of USP7i on BCOR, H2AK119ub1 and H3K27me3 is illustrated at the *HOXA* gene cluster, which represents a Polycomb chromatin domain (Figure 6D).

**Figure 6.**
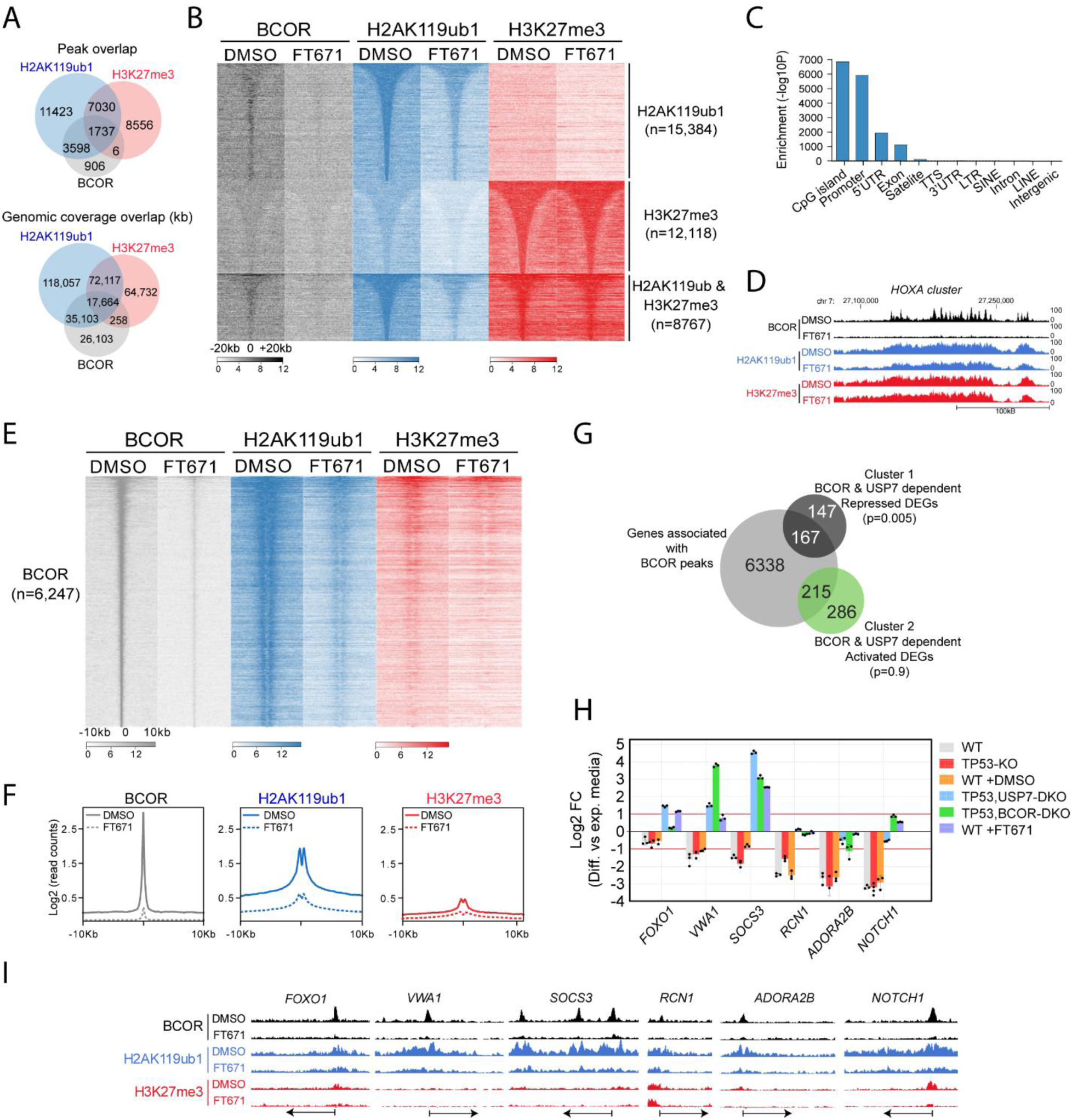
USP7 promotes H2AK119ub1 deposition via BCOR-PRC1.1 (A) Venn diagram showing the overlap between BCOR, H2AK119ub1 and H3K27me3 ChIP-seq peaks in control SH-SY5Y cells grown in differentiation medium for 3 days supplemented with DMSO. (B) Heatmap showing BCOR, H2AK119ub1 and H3K27me3 signals in SH-SY5Y cells grown in differentiation medium for 3 days supplemented with either DMSO or FT671. Average ChIP-seq read counts of two biological replicates per condition are shown in 50 bp bins and in 40 kb windows. (C) Barplot showing enrichment (-Log10 false discovery rate) of genomic features of BCOR binding sites. (D) Genome browser view of BCOR, H2AK119ub1 and H3K27me3 signals at the *HoxA* cluster. (E) Heatmap showing BCOR, H2AK119ub1 and H3K27me3 signals at all BCOR binding sites in SH-SY5Y cells grown in neural differentiation medium for 3 days (in the presence or absence of FT671). Average ChIP-seq read counts of two biological replicates per condition are shown in 50 bp bins and in 20 kb windows. (F) Average plot quantification of BCOR, H2AK119ub1 and H3K27me3 mean signals across BCOR binding sites shown in (E). (G) Venn diagram showing the overlap between genes associated with BCOR loci and BCOR- and USP7-dependent DEGs associated with neural differentiation, identified in Figure 5F. Cluster 1: USP7 and BCOR dependent genes that are repressed during neuronal differentiation; Cluster 2: USP7 and BCOR dependent genes that are activated during neuronal differentiation. Hypergeometric testing revealed that USP7- and BCOR-dependent repressed genes are significantly enriched for BCOR binding (p=0.005) while activated genes are not (p=0.9). (H) Barplot showing examples of mRNA expression of direct target genes of BCOR that are USP7-dependent. Means and SDs were derived from 3 biological replicates. For RNA-seq data see Data S3. (I) UCSC genome browser views of BCOR, H2AK119ub1 and H3K27me3 signals at representative loci. Each track represents the merged read counts of two biological replicates.

Next, we examined the relationship between chromatin state and gene expression at loci bound by BCOR. USP7i eliminates BCOR binding to chromatin, which is accompanied by a substantial loss of H2AK119ub1 and a modest reduction of the already low H3K27me3 mark (Figures 6E-F). We assigned BCOR peaks to nearby genes (n=6,720) and intersected those with BCOR and USP7 dependent DEGs that are associated with neural differentiation (identified in Figure 5F). Notably, genes that are repressed by BCOR and USP7 during neural differentiation (Figure 6G, cluster 1), are significantly enriched for BCOR binding (p=0.005). In contrast, genes that are activated in a BCOR and USP7 dependent manner (cluster 2) are not significantly enriched for BCOR binding (p=0.90). Genes repressed by USP7 and BCOR include key (neuro)developmental regulators, such as *FOXO1*, *VWA1*, *RCN1*, *ADORA2B* and *NOTCH1* (Figure 6H). Genome browser views illustrate the loss of BCOR and H2AK119ub1 at these genes in the presence of FT671 (Figure 6I). Collectively, our results imply that BCOR-ncPRC1.1 is a major downstream effector of USP7 in the regulation of neurodevelopmental gene expression.

## DISCUSSION

To gain insight into the molecular basis of Hao-Fountain syndrome, we have begun to uncover the substrates and pathways relevant for neuronal differentiation. We analyzed the transcriptomic, proteomic and epigenomic impact of chemical inhibition and genetic ablation of USP7 in cell models of neuronal differentiation. We established that USP7 activity is required for differentiation of ESCs to NSCs, for neuronal differentiation of i3N-iPSCs and SH-SY5Y cells, and for axon guidance in *Drosophila*. Our multi-omics analysis provides detailed insights into the USP7- dependent protein and transcriptional circuitry underpinning neurodifferentiation. We established that, out of a multitude of USP7 substrates (Figure 3L), BCOR-ncPRC1.1 mediates a large portion of USP7-dependent gene regulation that underpins neuronal differentiation (Figure 7). USP7 activity is essential for the integrity of BCOR-ncPRC1.1 and its binding to the promoter regions of target genes, where it deposits H2AK119ub1. In agreement with the crucial role of BCOR- ncPRC1.1 in the transcriptional control of neuronal differentiation, loss-of-function mutations in multiple signature subunits, including BCOR, RING1A/B and KDM2B, have been implicated in distinct NDDs. Our results suggest that ncPRC1.1-associated NDDs and Hao-Fountain syndrome involve dysregulation of a shared epigenetic network that is crucial for neuronal cell differentiation.

**Figure 7.**
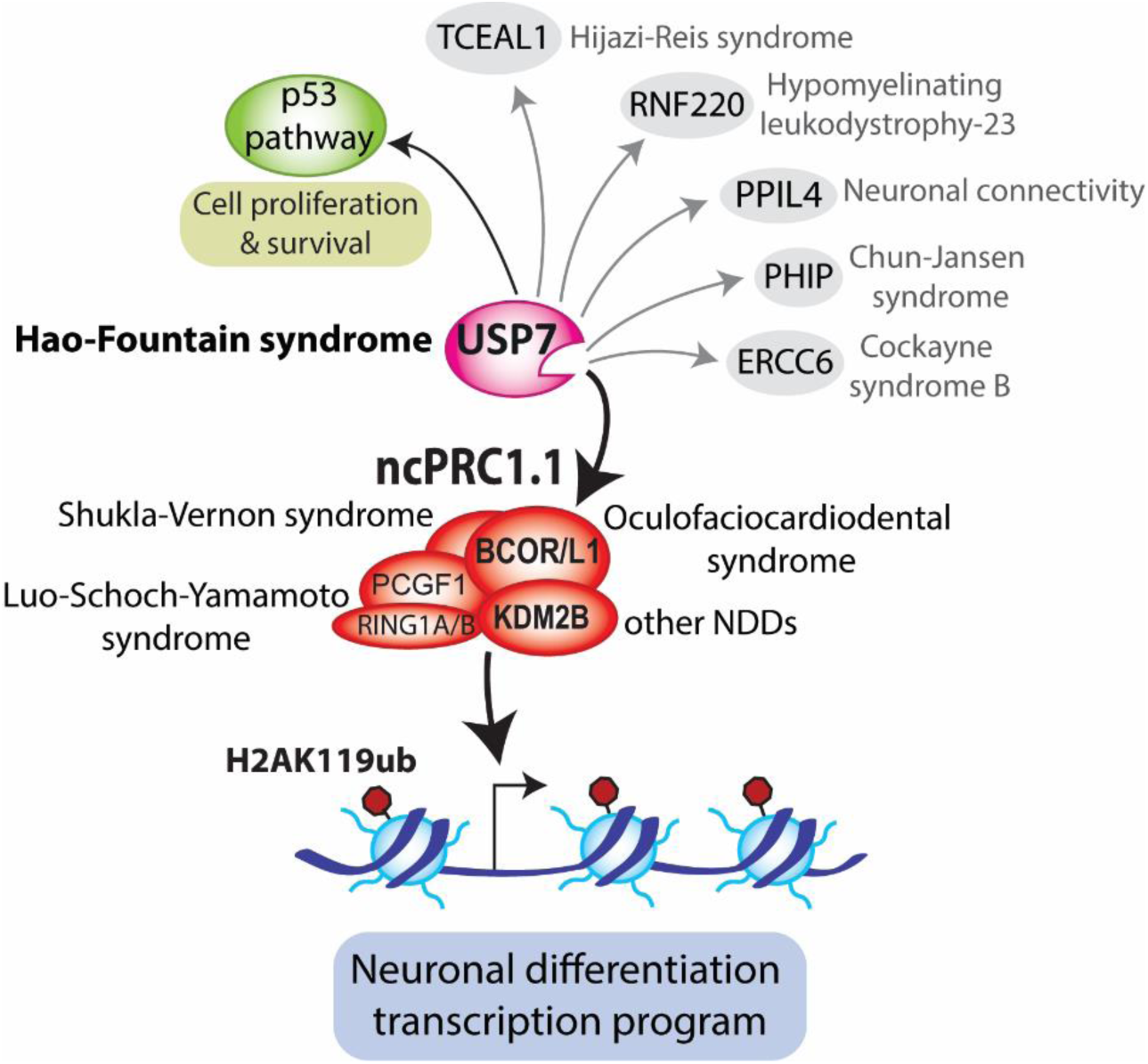
USP7 regulates neuronal differentiation via BCOR-PRC1.1 Summary of results. BCOR-ncPRC1.1 mediates the majority of the USP7-dependent neurodifferentiation transcription program. USP7 activity is essential for BCOR-ncPRC1.1 stabilization and binding to promoter regions of neurodevelopmental genes, where it deposits H2AK119ub1. Loss-of-function mutations in different ncPRC1.1 subunits have been implicated in various NDDs. Thus, Hao-Fountain syndrome and ncPRC1.1-associated NDDs involve dysregulation of a common epigenetic network. Although USP7 also regulates ncPRC1.6, this Polycomb complex is not required for neuronal differentiation. Likewise, our results do not support a role in neuronal differentiation for another well-established USP7 substrate, TRIM27. However, p53-directed regulation of cell proliferation and survival is likely to affect Hao-Fountain syndrome. Lastly, we and others identified additional USP7 substrates that are associated with various NDDs. Dysregulation of these proteins due to reduced USP7 function is likely to contribute to Hao-Fountain syndrome. See main text for details and references.

While USP7 binds and stabilizes both ncPRC1.1 and ncPRC1.6, our results indicate that only ncPRC1.1 is essential for the USP7-dependent program of neuronal differentiation. These results reflect the functional diversification between distinct PRCs.^20,21^ Indeed, there is a preponderance of mutations in ncPRC1.1, but not in ncPRC1.6, that cause NDDs. Pathogenic variants in the X-linked gene *BCOR* cause oculofaciocardiodental (OFCD) syndrome (OMIM #300166), which is a dominant disorder that affects females and is characterized by ocular, dental, cardiac and skeletal anomalies and mental retardation.^39,40^ Hemizygous *BCOR* loss-of-function in males causes gestational lethality, while hypomorphic missense mutations cause a severe microphthalmia syndrome. Pathogenic variants in the BCOR paralog *BCORL1* cause Shukla- Vernon syndrome (OMIM #301029), characterized by intellectual disability, behavioral difficulties, and dysmorphic features.^41^ Heterozygous loss of *KDM2B* function has been associated with developmental delay, intellectual disability, autism, anomalies of the eyes, heart and urogenital system.^42^ While PCGF1 has not been implicated in a human NDD, PCGF1-deficient mice display an array of neurodevelopmental phenotypes (www.mousephenotype.org/data/genes/MGI:1917087). Missense mutations in RING1A and RING1B (OMIM 619460) that impair deposition of H2AK119ub1 have also been implicated in NDDs.^43,44^ Pertinently, the link between loss of H2AK119ub1 and NDDs supports the relevance of this Polycomb-associated histone mark in human developmental gene regulation.

The various ncPRC1.1-associated NDDs and Hao-Fountain syndrome present overlapping, but also unique symptoms, reflecting shared and distinctive functionalities of different ncPRC1.1 subunits and USP7. Dissection of ncPRC1.1 in human ESCs revealed that BCOR, but not KDM2B, is crucial for the maintenance of primed pluripotency.^45^ In mouse ESCs, however, BCOR appears to be dispensable for pluripotency.^46^ Moreover, it was shown that the recruitment of BCOR-ncPRC1.1 to a subset of crucial target genes was independent of KDM2B, and that BCOR can repress transcription independently of H2AK119ub1.^45^ Our previous proteomics analysis revealed surprisingly large differences in relative MS abundance scores of individual ncPRC1.1 components upon immunopurification via different subunits.^10^ These results indicate that ncPRC1.1 might represent more a dynamic protein association rather than a static stoichiometric complex. Collectively, these observations suggest that ncPRC1.1 does not behave as a monolithic entity, but that different subunits can provide independent functionalities.

We also examined the consequences of missense and truncating variants in USP7, associated with Hao-Fountain syndrome.^2^ Some variants of USP7 were expressed at a lower level or were mislocalized in the cytoplasm, while others had an impaired ability to stabilize substrates. Although a more extensive analysis is required, our results indicate that different substrates display different sensitivities to some of the USP7 variants. Particularly BCOR seems to be more sensitive to some missense mutations than other substrates. Moreover, USP7-M225I and USP7-G392D might be more active than wt USP7 in the stabilization of some substrates. Thus, Hao-Fountain syndrome might not only be caused by a loss of USP7 function, but also by hyperactive gain-of-function variants.

H2AK119ub1 and H3K27me3 are the canonical chromatin marks associated with Polycomb repression. H2AK119ub1 and H3K27me3 co-exist at about a third of their loci in differentiating SH-SY5Y cells, but they are not globally coincident. These observations confirm that these marks can be deposited independent of each other at the majority of loci.^10,19,25,28^ Note, however, that canonical Polycomb targets, like the *HOX* clusters, are marked by both H2AK119ub1 and H3K27me3. Genomic loci bound by BCOR are predominantly characterized by H2AK119ub1, either in the presence or absence of H3K27me3. BCOR binds directly to about half of the BCOR- and USP7-dependent genes that are associated with neural differentiation. Loss of USP7 activity abrogated chromatin binding of BCOR and caused a substantial drop in the levels of genomic H2AK119ub1, with only modest effects on H3K27me3 (see also (10)). We note that the global reduction in H2AK119ub1 is caused by the loss of both ncPRC1.1 and ncPRC1.6, as well as potential indirect effects. Emphasizing the importance of H2AK119ub1 dosage, loss of function mutations in BAP1, the enzyme that deubiquitylates H2AK119ub1, cause the NDD Brainbridge- Ropers syndrome (OMIM **#** 615485;(47)). Collectively, these observations suggest that developmental gene regulation requires the maintenance of H2AK119ub1 levels within an optimal window.^48^

The results presented here emphasize a major role for BCOR-ncPRC1.1 as a, p53- independent, mediator of the USP7-controlled neurodevelopmental transcription program. However, given USP7’s plethora of substrates, additional pathways are likely to contribute to Hao-Fountain syndrome (Figure 7). Studies in mice indicated that loss of USP7 also affects (neuro)development via p53-directed control of cell proliferation and survival.^14,15^, Another function of USP7 that has been linked to Hao-Fountain syndrome is the regulation of endosomal recycling through stabilization of a MAGEL2-TRIM27 complex.^1^ However, our results do not support a role for TRIM27 in neuronal differentiation (Figure 5A). Indeed, the predominantly nuclear localization of USP7 is difficult to reconcile with an endosomal function. Recently, USP7 was implicated in regulation of neuronal connectivity in the mouse brain via splicing factor PPIL4.^49^ In addition to BCOR-ncPRC1.1, USP7 has other targets that have been associated with human NDDs, including e.g., ERCC6 (Cockayne syndrome B, OMIM #133540; ^(50)^), RNF220 (Hypomyelinating leukodystrophy-23, OMIM #619688; ^(51)^), TCEAL1 (Hijazi-Reis syndrome, OMIM #301094; ^(52,53)^), and PHIP/BRWD2 (Chung-Jansen syndrome, OMIM #617991; ^(54)^).

Indeed, loss of PHIP also impaired neuronal differentiation of SH-SY5Y cells (Figure 5A). Thus, the complex pathology of Hao-Fountain syndrome is likely to be the result of the cumulative effects of reduced functioning of a subset of USP7 substrates (Figure 7). Therefore, it is imperative to investigate the contribution of additional USP7 substrates to Hao-Fountain syndrome and other NDDs. Our identification of BCOR-ncPRC1.1 as a key downstream target of USP7 during neurodifferentiation provides a template for such future studies.

## MATERIALS & METHODS

### Cell culture and neuronal differentiation

See Table S1 for an overview of reagents. Cells were maintained at 37°C in a humidified incubator with 5% CO2 and routinely checked for mycoplasma using the MycoAlert kit. Unless indicated otherwise, cell cultures were split at ∼80% confluency. To split cells, media was aspirated, cells were washed with PBS (137 mM NaCl, 2.7 mM KCl, 10 mM Na2HPO4 and 1.8 mM KH2PO4, pH7.4) at 37°C, which was replaced with PBS containing 0.05% Trypsin and 0.53 mM EDTA. Following cell detachment, fresh media with fetal bovine serum (FBS) was added, cells were resuspended and plated. HEK293T cells were cultured in DMEM containing 10% FBS and 1x Pen/Strep (100 units/ml Penicillin and 100 µg/ml Streptomycin). H9 ESCs were cultured in mTeSR medium containing 1x mTeSR supplement and 1x Pen/Strep on 6-wells plates coated with Matrigel (diluted 1:100 in DMEM:Nutrient Mixture F-12 (DMEM-F12)). H9 cells were detached with TrypLE. After collection, cells were resuspended in 2 ml mTeSR containing 0.01 µM ROCK-inhibitor and seeded. Following cell attachment, medium was replaced by mTeSR without ROCK-inhibitor. For differentiation to NSCs, H9 cells were seeded on Matrigel-coated 12-wells plates at a density of ∼1.8x10^4 cells/cm^2^ in mTeSR. At ∼90% confluency, cells were washed with PBS, followed by the addition of Differentiation medium (1x Knockout DMEM, 15% Knockout Serum Replacement, 1x L-glutamine, 1x Non-Essential Amino Acids (NEAA), 50 µM ß- mercaptoethanol, 1x Pen/Strep, supplemented with dual SMAD inhibitors: 2 µM A83-10 and 2µM Dorsomorphine). For USP7i experiments, H9 cells were grown in the presence or absence of 10 µM FT671 or its solvent 7 mM DMSO, as a control. For the next five days, the medium was refreshed daily. At day six, the medium was replaced with a 1:1 mix of Differentiation medium and NSC medium (1x Knockout DMEM-F12, 1x L-glutamine, 20 ng/ml EGF, 5 ng/ml bFGF, 2% StemPro Neural Supplement, 1x Pen/Strep, 2µM A83-10 and 2µM Dorsomorphine). Medium was refreshed every other day for 4 days. Thereafter, cells could be expanded by splitting, for which 250 µl Accutase was used to detach the cells, which were cultured in NSC medium. Maintenance, induction and maturation of i3N iPSCs was as described.^55^ Briefly, i3N iPSCs were maintained in StemFlex medium using Geltrex coated culture plastics. After seeding ∼2x10^5 cells per well in a 6-wells plate, neuronal differentiation was induced by addition of 2 μg/ml doxycycline. For neuron maturation, medium was replaced by Cortical Neuron Culture (CNC) medium (Neurobasal medium supplemented with 1x B27 supplement, 10 ng/ml BDNF and 10 ng/ml NT3). For USP7i, maturation in CNC medium was in the presence or absence of 10 µM FT671 or its solvent 7 mM DMSO, as a control. SH-SY5Y cells were cultured in DMEM-F12 containing 15% FBS and 1x Pen/Strep and split at ∼60-70% confluency. Neuritogenesis was induced by culturing the cells in differentiation medium (Neurobasal plus medium supplemented with 1x B27, 2 mM Ultraglutamine (Glutamax) and 10 µM Retinoic Acid), in the dark for 10 days. Medium was refreshed every 3-4 days. To generate neurospheres, ∼8x10^4 cells per well were seeded in non- adherend 96-wells U bottom-plates, at a final volume of 100 μl, and centrifuged for 3 min. at 1,200 rpm. Next day, an additional 100 μl fresh medium was added. After 4 days, medium was removed, spheres were washed 1x with PBS and next cultured in differentiation medium in the dark. Medium was refreshed at day 3-4. After 7 days, spheroids were transferred to a 1.5 ml Eppendorf tube, medium removed and replaced with 40 μl ice-cold mixture of 2/3 Basement Membrane Extract (BME) and 1/3 differentiation medium. Next, spheroids were transferred to a droplet of BME- differentiation medium in a pre-warmed 48-wells plate. BME was allowed to solidify by putting the plate upside down in a 37°C incubator for 40 min. Neurospheres were cultured for three more days in 250 μl differentiation medium. *Drosophila* S2 cells were cultured in Schneider’s *Drosophila* medium supplemented with 10% FBS and 1x Pen/Strep at 25C under normal atmospheric conditions. Cell viability was determined using the CellTiter-Glo® 2.0 assay (Promega). In short, ∼5000 cells/well were seeded in a 96 well plate. Next day, medium was replaced with either expansion or differentiation medium, containing FT671 (10 µM) or DMSO, as indicated. For each condition three biological replicates were used. After 1 (H9) or 4 days (SH- SY5Y) cells were lysed by adding CellTiter-Glo 2.0 reagent (1:1) to the wells. Lysates were transferred to an opaque 96 well plate and luminescence was measured in a Glomax™ 96 microplate luminometer (Promega). To compare SH-SY5Y p53-KO cells with SH-SY5Y p53, USP7-DKO cells in differentiation medium, the luminescence was normalized against the luminescence of these cells in serum-free expansion medium.

### Generation of cell lines

Guides against *USP7*, *TP53*, *BCOR*, *PCGF6*, *PHIP, RING1A*, *RING1B*, and *TRIM27* (see table S1) were cloned into the Esp3I restriction site of pLentiCRISPRv2GFP. The integrity of all constructs was verified by DNA sequencing. KO cell lines were generated by CRISPR-Cas9- mediated genome editing. For p53-KO cells, PX459 guide constructs were transfected (1.25 µg DNA each, per well of a 6-wells plate) into SH-SY5Y cells using FuGENE HD Transfection Reagent. After 24 hours, cells were selected with puromycin (1 µg/ml) for three days. Surviving cells were diluted and plated on 10 cm dishes for colony formation. For the other KO lines, guide vectors were transfected into wt or p53-KO#F12 SH-SY5Y cells or into HEK293T cells (2.5 μg DNA per well of a 6-wells plate). 48 hrs after transfection, SH-SY5Y cells expressing GFP were FACS sorted as a pool of ∼4,000 cells per well of a 48-wells plate, containing filtered conditioned (40%) medium from the parental cell line, and cultivated for 3 days. Next, cells were split into 10 cm petridishes, containing 10% conditioned medium. 12-15 days later, colonies were picked and grown in 96-wells plates. HEK293T cells expressing GFP were FACS sorted as single cells in 96- wells plates and screened. The p53,USP7-DKO#41 line was derived from p53-KO#D9 using the USP7 sgRNA CRISPR Lentivector set. Clones were cultured, duplicated and screened by immunoblotting (for antibodies see key resources table). To produce the SHSY5Y USP7^het^ line, plasmids containing guide USP7_HGLibA_53409 I and USP7 HGLibA_53411 III were transfected, GFP-positive cells were FACS sorted, cultivated, and monoclonals were picked as described above. Genomic DNA was isolated from clones using the Quick extract buffer, according to the manufacurers protocol, and screened by PCR. Selected p53-KO, p53,BCOR- DKO, p53,USP7-DKO, USP7^het^ and TRIM27-KO lines were verified by mass spectrometry.

### Drosophila procedures

*Drosophila* lines were obtained from the Bloomington *Drosophila* Stock Center (https://bdsc.indiana.edu). The *Usp7^KG068^*^14^ mutant has been described.^13^ Stocks were maintained on standard corn medium at 18°C. All crosses and experiments were carried out at room temperature (RT). 3^rd^ instar larval brains, with attached eye imaginal discs, were dissected in PBST (PBS, 0.1% Triton-X100), fixated with 4% CH2O for 20 min at RT, washed gently with PBST and then blocked with PBST containing 5% dry milk and 5% BSA for 2hrs at RT. Following 3 washes with PBST, brains were incubated with α-CHAOPTIN (mouse 24B10) and α-ELAV (rat 7E8A10) monoclonal antibodies (Developmental Studies Hybridoma Bank) diluted 1:200 in PBST/2%BSA overnight (ON) at 4°C. Following 3 washes with PBST, brains were incubated with secondary antibodies (Alexa Fluor 594 α-mouse and Alexa fluor 488 α-rat) diluted 1:200 in PBST for 2 hrs at RT. Brains were washed 3x with PBST and mounted in Vectashield containing DAPI.

### Imaging

For phase contrast imaging, 1.4x10^5 cells were seeded in 35 mm dishes. For IF, 2.3x10^4 cells were split on 18 mm sterilized coverslip in a 12 wells plate one day before the addition of differentiation medium. For IF of i3N iPSCs, cells were grown on coverslips that were surface-activated by plasma treatment for 1-2 min using a Laboratory Corona Treater, MODEL BD-20 (Electro-Technic Products), followed by coating with 50 µg/ml poly-D-lysine in PBS for 20 min at 37°C. All washes and incubations were on an orbital shaker. The coverslips were washed several times with PBS and then coated with murine laminin (1.5 µg/ml) in PBS for 20 min at 37°C and, finally, washed with PBS before use. For IF, cells grown on coverslips were washed 3x with PBS at RT, fixed with ice-cold 100% methanol for 15 min on ice, and washed 3x with PBS for 5 min on an orbital shaker. Next, coverslips were blocked using filtered 1% BSA in PBS/0.3%Triton X- 100 for 60 min at RT on a shaker. The coverslips were incubated with α-βTubIII (abcam, 1:1200), α-USP7 (abcam, ab101648, 1:600), α-αTubulin (Sigma, T5168, 1:750) or α-Flag (Sigma, F3165, 1:1800) primary antibodies diluted in PBS/0.3%TritonX-100/1%BSA were incubated ON at 4°C under gentle shaking. Next, coverslips were washed 3x with PBS and 3x with PBST (each wash for 5 min at RT), followed by incubation with Alexa Fluor 488 α-mouse, Alexa Fluor 594 α-mouse or Alexa Fluor 488 α-rabbit secondary antibodies diluted 1:1000 in PBS/1% BSA/0.05%Tween20 for 1 hr at RT in the dark. Next, coverslips were washed 3x with PBS/0.05%Tween 20, 1x with PBS and 3x with demineralized H2O. Coverslips were mounted in Vectashield, containing DAPI, and stored at 4°C in the dark. Spheroids were washed 3x with PBS, followed by fixation with 750 μl 4% CH2O in PBS at RT for 30 min. Following fixation, spheroids were washed 3x with PBS, incubated in PBS/0.1M glycine for 30 min at RT, followed by 1x wash with PBS. For permeabilization, the spheroids were incubated with PBS/0.5%TritonX-100 for 30 min at RT, washed 1x with PBS/0.5%FBS and then blocked for 2 hrs at RT in 750μl PBSTDF (PBS, 0.3% Triton X-100, 1% DMSO, 0.5% FBS). Spheroids were incubated with α-βTubIII antibodies (1:1200) in PBSTDF ON at 4°C. The next day, spheroids were washed 5x with PBS/0.5%FBS for 10 min at RT. For the following steps, plates were kept in the dark. Incubation with secondary antibodies was in PBSTDF for 2 hrs at RT, followed by 5 washes for 10 min at RT with PBS/0.5%FBS. 20 μl mounting media with DAPI was added to each well and incubated for 1 hour at RT. Plates were stored at 4°C in the dark. Phase contrast images were acquired using a Olympus IX 70 microscope equipped with an 10x UPlanFI 0.30 Ph1 objective. Images were processed and the brightness/contrast was adjusted using Fiji/ImageJ software (https://imagej.net). Phase contrast images were used to quantify the percentage of differentiated cells, defined as having an extension that is at least 2x the length of the cell body, using Fiji/ImageJ software and the cell counter tool. Confocal images were acquired using a Leica TCS SP5 confocal microscope equipped with Argon 488 nm and HeNe 594 nm laser lines, a UV-lamp, and a 40x HCX PL APO CS 1.3 NA oil-immersion objective and Leica LAS-AF acquisition software. Images were analyzed and processed using Fiji/ImageJ software.

### Co-transfection experiments and immunoblotting

Experiments were performed essentially as described.^10^ Briefly, HEK293T USP7-KO cells were co-transfected with vectors expressing GFP-tagged substrates in combination with either an empty vector or a vector expressing 3xFlag-USP7 wt and variants (see key resources table) and a vector expressing His-tagged ubiquitin. Cells were seeded in 12-wells plates. The next day, medium was replaced with serum-free DMEM, and transfected with 1.5-2.5 µg of total DNA mixed with 4.5-7.5 µg of polyethylenimine in a total volume of 200 µl PBS. When indicated, FT671 (10 µM) was added 24 hours before harvesting. Next day, the medium was replaced with DMEM/10%FBS. After 48 hrs, medium was removed by aspiration, cells were washed 1x with PBS, and then were lysed on the plate by the addition of 1x Laemmli buffer (63 mM Tris-HCl pH 6.8, 2% SDS, 10% Glycerol, 0.0005% Bromophenol blue, containing 0.1M DTT), and transferred to a 1.5ml Eppendorf tube. Samples were sonicated for 5 min on a 30 sec on/off cycle in a Bioruptor® UCD- 200 sonicator and heated for 3 minutes at 95°C. Samples were analyzed by immunoblotting as described.^10^

### Immunopurification of endogenous USP7

Wt and p53,USP7-DKO#58 SH-SY5Y cells were harvested from one 10 cm plate per triplicate, following 3 washes with PBS, by scraping in 10 ml of PBS. All following procedures were on ice or at 4°C. Incubations were on a rotating wheel. Cells were collected in 15 ml falcon centrifuge tubes and centrifuged at 1,000 g for 5 min at 4°C. Following aspiration, cell pellets were resuspended in 750 μl HEMG/150 buffer (25 mM HEPES-KOH pH7.6, 5 mM EDTA, 1.5 mM MgCl2, 10% glycerol, 150 mM NaCl, 0.1% Nonidet P40 and protease inhibitors: 1 μM aprotinin, 1 μM leupeptin, 1 μM pepstatin) and sonicated for 5 min using a 30 sec on/off cycle. Next, lysates were incubated with benzonase (100 U/ml) for 30 min. After addition of ethidium bromide (50 μg/ml), lysates were cleared by centrifugation at 20,000 g for 15 min, and the supernatants were transferred to 1.5 ml eppendorf tubes. Cleared lysates were incubated with 7.5 μl α-USP7 (Bethyl Laboratories, A300-033A) antibodies per lysate for 2.5 hrs. Next, 50 μl of protein-A-Sepharose beads slurry in HEMG/150 (1:5 ratio of beads:buffer) was added and incubated for 2 hrs. Following centrifugation for 3 min at 3,000 rpm, beads were washed 3x with 1 ml HEG/150/0.1%Nonidet P40 (25 mM HEPES-KOH pH7.6, 1.5 mM EDTA, 10% glycerol, 150 mM NaCl, 0.1% Nonidet P40 and protease inhibitors), 3x with HEG/150/0.01%Nonidet P40, and finally 2x with HEG/100 without Nonidet P40. The resulting immunopurified proteins on beads were directly processed for mass spectrometric analysis.

### Proteomics and mass spectrometry

For IP-MS, proteins were digested on-bead with sequencing grade trypsin (1:100 (w:w), Roche) ON at RT. Protein digests were desalted using a C18 Stagetip (2 plugs of 3M Empore C18) and eluted with 80 % acetonitrile and dried in a Speedvac centrifuge. Peptides were then analyzed by nanoflow LC-MS/MS, as described below. For global proteome analysis of whole cell extracts, cells were lysed in 100 mM Tris/HCl, pH 8.2, containing 1% sodium deoxycholate (SDC) using sonication in a Bioruptor Pico (Diagenode). Protein concentrations were measured using the BCA assay (ThermoFisher Scientific). 100 μg protein was reduced in lysis buffer with 5 mM dithiothreitol and alkylated with 10 mM iodoacetamide. Next, proteins were digested with 2.5 μg trypsin (1:40 enzyme:substrate ratio) ON at 37 °C. After digestion, peptides were acidified with trifluoroacetic acid (TFA) to a final concentration of 0.5 % and centrifuged at 10,000 g for 10 min to spin down the precipitated SDC. Peptides in the supernatant were desalted on a 50 mg C18 Sep-Pak Vac cartridge (Waters). After washing the cartridge with 0.1 % TFA, peptides were eluted with 50 % acetonitrile and dried in a Speedvac centrifuge. Peptides were then analyzed by nanoflow LC-MS/MS. Nanoflow LC-MS/MS was performed on an EASY-nLC system (Thermo) coupled to an Orbitrap Lumos or Eclipse Tribrid mass spectrometer (both ThermoFisher Scientific) or on a Vanquish Neo LC system (Thermo) coupled to an Orbitrap Exploris 480 (Thermo), all operating in positive mode and equipped with a nanospray source. Peptide mixtures were trapped on a PepMap trapping column (2 cm × 100 µm, Thermo, 164750) at a flow rate of 1 µl/min. Peptide separation was performed on ReproSil C18 reversed phase column (Dr Maisch GmbH; column dimensions 25 cm × 75 µm, packed in-house) using a linear gradient from 0 to 80% B (A = 0.1% FA; B = 80% (v/v) AcN, 0.1 % FA) in 120 min and at a constant flow rate of 250 nl/min. The column eluent was directly sprayed into the ESI source of the mass spectrometer. For data dependent acquisition (DDA), all mass spectra were acquired in profile mode and the resolution in MS1 mode was set to 120,000 (automatic gain control (AGC) target: 4E5) and the m/z range to 350-1400. Fragmentation of precursors was performed in 2 s cycle time data-dependent mode by higher-energy collisional dissociation (HCD, or beam-type collision induced dissociation (CID)) with a precursor window of 1.6 m/z and a normalized collision energy of 30.0; MS2 spectra were recorded in the orbitrap at 30,000 resolution. Singly charged precursors were excluded from fragmentation and the dynamic exclusion was set to 60 seconds. For data independent acquisition (DIA), all spectra were recorded at a resolution of 120,000 for full scans in a 350–1100 m/z scan range. The maximum injection time was set to 50 ms (AGC target: 4E5). For MS2 acquisition, the mass range was set to 336–1391 m/z with variable isolation windows ranging from 7–82 m/z with a window overlap of 1 m/z. The orbitrap resolution for MS2 scans was set to 30,000. The maximum injection time was at 54 ms (AGC target: 5E4; normalized AGC target: 100 %). For targeted proteomics, a PRM regime was used to select for a set of previously selected peptides on an Orbitrap Eclipse Tribrid or an Orbitrap Exploris 480 mass spectrometer operating in positive mode. Precursors were selected in the quadrupole with an isolation width of 0.7 m/z and fragmented with HCD using 30 % collision energy (CE). MS1 and MS2 spectra were recorded in the orbitrap at 30,000 resolution in profile mode and with standard AGC target settings. The injection time mode was set to dynamic with a minimum of 9 points across the peak. The sequence of sampling was blanks first and then in order of increasing peptide input amounts to avoid any contamination of previous samples.

### Mass spectrometry data analysis

DDA raw data files were analyzed using the MaxQuant software suite (version 2.2.0.0, www.maxquant.org)^56^ for the identification and relative quantification of proteins. ‘Match between runs’ was disabled and a false discovery rate (FDR) of 0.01 for peptides and proteins and a minimum peptide length of 6 amino acids were required. The Andromeda search engine was used to search the MS/MS spectra against the *Homo sapiens* Uniprot database (version May 2022) concatenated with the reversed versions of all sequences and a contaminant database listing typical background proteins. A maximum of two missed cleavages were allowed. MS/MS spectra were analyzed using MaxQuant’s default settings for Orbitrap and ion trap spectra. The maximum precursor ion charge state used for searching was 7 and the enzyme specificity was set to trypsin. Further modifications were cysteine carbamidomethylation (fixed) as well as methionine oxidation (variable). The minimum number of peptides for positive protein identification was set to 2. The minimum number of razor and unique peptides set to 1. Only unique and razor non-modified, methionine oxidized and protein N-terminal acetylated peptides were used for protein quantitation. The minimal score for modified peptides was set to 40 (default value). DIA raw data files were analyzed with the Spectronaut Pulsar X software package (version 17.0.221202, www.biognosys.com,), using directDIA for DIA analysis including MaxLFQ as the LFQ method and Spectronaut’s IDPicker algorithm for protein inference. The Q-value cutoff at precursor and protein level was set to 0.01. All imputation of missing values was disabled. PRM data were analyzed with Skyline (version 23.0.1.268, <u>skyline.ms</u>). Skyline reports containing all essential data such as area per fragment for each target were loaded into R and curated manually. If the expected peptide fragment ion peaks could not be correctly assigned, fragments were excluded and their associated areas were set to 0. Summed areas were calculated for all fragment ion peak chromatograms for each target peptide and then summed over all targeted peptides per protein. Standard deviations of at least three biological replicates were calculated for every target peptide in each condition and indicated in the bar plots. All summed areas were normalized to the WT condition. Peptide bar graphs were constructed by dividing the bar value for each target peptide into the respective areas for every fragment ion peak. For the protein abundance comparison between non-differentiated WT and other conditions, the normalized areas for all detected fragment ion chromatograms were summed over all selected target peptides per protein and visualized in box plots. The median for the WT condition was set to 1. All protein abundances are based on ≥ 2 target tryptic peptides unless indicated otherwise. Downstream data analysis was performed using Perseus (www.maxquant.org/perseus/), Prism (v 10, www.graphpad.com) or in-house developed software tools. In Perseus, data from MaxQuant or Spectronaut searches were imported and all peptide and protein intensity data were first LOG2 transformed. For PCA, protein entries were required to have valid intensity values for all conditions and replicates. For the protein profile plots, normalization by Z score transformation was performed, where the mean of each row was subtracted from each value in that row. The result was then divided by the standard deviation of that row. Z scored values were used as the input for fuzzy C means clustering to generate the profile plots from which populations of proteins with similar dynamic abundance profiles were selected and categorized. The top 100 proteins that resembled the reference profile most based on the lowest Pearson distances were selected for further gene ontology analysis. Both for the USP7- KO and USP7i experiments, six distinct categories were defined, i.e. (1, blue) proteins upregulated in WT, but not in USP7-KO (c.q. USP7i) after differentiation only, (2, pink) proteins downregulated in the in WT, but not in USP7-KO (c.q. USP7i) upon differentiation only, (3, yellow) proteins downregulated in USP7-KO (c.q. USP7i) independent of differentiation status, (4, green) proteins upregulated in USP7-KO (c.q. USP7i) independent of differentiation status, (5, orange) proteins upregulated during differentiation in a USP7 independent fashion, and, (6, brown) proteins downregulated during differentiation in a USP7 independent fashion. GO analysis was performed with g:Profiler (https://biit.cs.ut.ee/gprofiler/gost). Protein accession ID’s were uploaded to the g:GOSt tool, which performs functional enrichment analysis, a.k.a. over- representation analysis (ORA). Proteins were mapped to known functional information sources and statistically significantly enriched terms were detected. Benjamini-Hochberg FDR was used to determine the significance threshold, while the user threshold was set to 0.05. For further statistical analysis in GraphPad Prism, the ANOVA-like non-parametric Friedman test was applied, including a correction for multiple comparisons using Dunn’s statistical hypothesis testing. MS raw data and data for protein identification and quantification were submitted as supplementary tables to the ProteomeXchange Consortium via the PRIDE partner repository with the data identifier PXD051616.

### RNA isolation, sequencing and analysis

Biological triplicates were grown on 12-wells plates (H9 ESCs, 2 wells per triplicate) or 10-cm plates (SH-SY5Y cells, 1 plate per triplicate). RNA was isolated using TRI Reagent (Sigma Aldrich), following the manufacturers protocol. Medium was removed from the cells and the TRI Reagent was added directly onto the cells (1 ml TRI Reagent per well of a 12-wells plate and 5 ml per 10-cm plate). 1 ml of TriPure lysate (corresponding to 2 wells, or 1/5^th^ of a 10-cm plate) was transferred to a 1.5 ml Eppendorf tube and incubated for 5 min at RT. Next, 0.2 ml chloroform was added, shaken vigorously, and left for 15 min at RT. Samples were centrifuged for 15 min at 12,000 g at 4°C. The aqueous phase was transferred to a fresh tube, 0.5 ml isopropanol was added, mixed and incubated for 10 min at RT, followed by centrifugation at 12,000 g at 4° for 10 min. The precipitated RNA pellet was washed 2x with 1 ml 75% ethanol, dried for 3 min in a SpeedVac and resuspended in 100 μl of nuclease-free H2O. RNA concentrations and purities were determined using a Nanodrop spectrophotometer. RNA samples were processed by BGI Tech Solutions (Hong Kong): mRNA molecules were purified from total RNA using oligo(dT)-attached magnetic beads, cDNA libraries prepared and sequenced on the DNBSEQ high-throughput platform. Resulting FASTQ files were aligned against the human reference genome (hg19) *STAR*(1) and default setting and reads were assigned to mRNAs using HTseq-count(2) with the following settings: -m union - s reverse -t exon. DESeq2(3) (bioconductor.org) was used for differential gene expression analysis. Genes were considered differentially expressed (DEGs) if they fulfilled the following criteria: normalized read counts ≥ 25, fold change ≥ 2, and FDR < 0.05 at one or more conditions. GO term enrichment analysis of DEGs was performed by gProfiler (https://biit.cs.ut.ee/gprofiler/gost), with KEGG, Reactome and WikiPathways data. We used the default settings of gProfiler, which includes all genes as background for GO Terms enrichment analyses. Only significantly enriched GO terms (FDR <0.05) were selected. To identify USP7 or BCOR-dependent genes, a neurodifferentiation signature was determined by selecting DEGs in differentiation medium versus expansion medium, in both wt and in p53-KO cells, yielding 2,178 genes. This list was used to establish the neurodifferentiation-associated DEGs in USP7i, p53,USP7-DKO or p53,BCOR- DKO cells. RNA-seq data has been deposited in the Gene Expression Omnibus (GEO) under the accession code GSE263492.

### ChIP-Rx-seq and analysis

Biological duplicates of SH-SY5Y cells were grown in two 15 cm dishes per condition, which were processed independently. Cells were grown for 3 days in differentiation medium in the presence of 10 µM FT671 or its solvent 7 mM DMSO. Cells were harvested by trypsinization, followed by cell counting. For crosslinking, ∼2x10^7 cells were resuspended in 10 ml PBS. After addition of CH2O (1%) cells were crosslinked for 10 min on a roller mixer. Crosslinking was quenched with 125 mM glycine for 5 min. Next, cells were washed twice with PBS and lysed in 1.8 ml lysis buffer (50 mM Tris pH 8.0, 10 mM EDTA, 1% SDS and protease inhibitors: 1 μg/ml aprotinin, 0.2 mM AEBSF, 1 μg/ml leupeptin and 1 μg/ml pepstatin A). Protease inhibitors were present at all subsequent steps, until elution. Crosslinked chromatin was fragmented to approximately 200 bp DNA length in a Bioruptor® Standard sonication system. Reference chromatin was prepared from ∼8x10^7 *Drosophila* S2 cells, as described above. SH-SY5Y chromatin (185 µl) was mixed with S2 chromatin (15 µl), diluted 10x in dilution buffer (20 mM Tris, pH 8.0, 2 mM EDTA, 150 mM NaCl and 1% Triton X-100) and cleared by centrifugation at 16,000 g for 25 min. For BCOR ChIP, the amounts of chromatin were doubled. After centrifugation, supernatants were transferred to fresh 2 ml tubes and incubated ON at 4°C with either 5 µl α-H2AK119ub1 (D27C4, Cell Signaling Technology #8240), 15 µl α-H3K27me3 (C36B11, Cell Signaling Technology, #9733) or 10 µl α-BCOR (E6V3R, Cell Signaling Technology, #63972) antibodies. Next, 50 µl of magnetic Dynabeads™ Protein A slurry (corresponding to 1.5 mg Dynabeads blocked ON in dilution buffer containing 1% BSA) was added per tube, and incubated for 3.5 hrs. Next, beads were washed 5x with wash buffer/150 mM NaCl (20 mM Tris, pH 8.0, 2 mM EDTA, 0.1% SDS, 1% Triton X-100, 150 mM NaCl) followed by a final wash step with wash buffer/500 mM NaCl. DNA was eluted with elution buffer (0.1 M NaHCO3, 1% SDS, 0.5 mg/ml proteinase K) for 2 hrs at 37°C, followed by ON incubation at 65°C. Eluted DNA was purified by 2x phenol/chloroform extraction, followed by ethanol precipitation. Libraries for sequencing were prepared using the NEBNext® Ultra™ II DNA Library Prep Kit for Illumina® (New England Biolabs, #E7645L) according to manufacturer’s instructions. AMPure® XP Beads (Beckman Coultier) were used for cleanup. Sequencing was performed by BGI Tech Solutions (Hong Kong) Co., Ltd. on an DNBSEQ-G400 sequencing platform. FASTQ files were mapped against the human genome (hg19) and *Drosophila* genome (Dm6) using Bowtie2 (bowtie-bio.sourceforge.net) and default parameters. SAM files were further filtered for uniquely mapped reads and high-quality mapping (-q 10) using SAMtools, and PCR duplicates were removed using MarkDuplicates in PICARD tools. Total reads in *Drosophila* BAM files were used to calculate the normalization factors as described.^57^ BAM files were then converted to BED files using BEDTools, and converted to BedGraph using the genomecov function of the BEDtools portfolio. BedGraphs were converted to BigWig files using bedGraphToBigWig from the UCSC tool suite for further visualisation using the Genome Browser, hosted by UC Santa Cruz. BCOR peaks were called using MACS2^58^ with default parameters, while H2AK119ub1 and H3K27me3 peaks were called by using SICER,^59^ with default settings and a window size of 200 bp and a gap size of 600 bp. Peaks overlapping with human blacklist regions were excluded using BEDTools. Peaks were merged whenever they were located within a 5 kb window to generate the final set of peaks. Reads counting within peaks was carried out on normalized BAM files using the intersect function from BEDTools. To assign ChIP peaks to corresponding promoters, the window function from BEDTools was applied to promoters within 1 kb distance to BCOR, H2AK119ub1 or H3K27me3 peaks. Promoters are determined as +/-2 kb of transcription start sites. Heatmaps were generated using Biofluff 3.0.4 (https://fluff.readthedocs.io) with the following parameters: -e 5000 -F 250 - m -S. For average plots, read counts were obtained via the EaSeq interactive software for analysis and visualization of ChIP-sequencing data (https://easeq.net/). Normalized BAM files were used to call peaks with the following parameters: unnormalised background, 20 kb window from and plotted using GraphPad Prism v10 (https://www.graphpad.com/). gProfiler (https://biit.cs.ut.ee/gprofiler/gost), was used for GO term enrichment analysis (FDR<0.05) of genes associated with BCOR, H2AK119ub1 or H3K27me3 peaks. We used the default settings of gProfiler, which includes all genes as background for GO Terms enrichment analyses. ChIP-Rx-seq data has been deposited in GEO under the accession code GSE263492.

## DATA AVAILABILITY

RNA-seq and ChIP-Rx-seq data have been deposited in the Gene Expression Omnibus (GEO) under the accession code GSE263492. The mass spectrometry proteomics data have been deposited to the ProteomeXchange Consortium via the PRIDE partner repository with the data set identifier PXD051616.

## ACKNOWLEDGEMENTS

We are grateful to our lab members, Jesper Svejstrup and Marc Timmers for helpful discussions. This work was supported in part by Dutch Research Council ECHO grant no. 711.014.001 to C.P.V., Medical Research Council New Investigator Research Grant no. MR/X008517/1, Royal Society Research Grant no. RG\R1\241024, and Leukaemia & Lymphoma NI (LLNI) research fund to Y.A., ZonMw Vidi, grant 09150172110002 to TSB, ZonMw PSIDER Doorbraakprojecten grant 10250042110005 and Brain and Behavior Research Foundation Young Investigator award, BBRF YI grant 30787 to K.L.

## AUTHOR CONTRIBUTIONS

Conceptualization: J.W.vdM. and C.P.V. initiated and designed the study with input from all authors. J.W.vdM., G.E.C. and J.vdK. conducted most of the experiments, with assistance from L.B., A.vdV., B.T. and A.S. E.K. performed the Drosophila experiments, A.L. and Y.A. performed the bio-informatic analysis of RNA-seq and ChIP-Rx-seq experiments, K.B., D.D., W.A.S.D., J.vdW.T. and J.D. performed mass spectrometry and proteomics data analysis, K.L. and T.S.B. assisted with the E.S.C. experiments, T.A. and J.vH. assisted with the iPSC experiments, J.W.vdM., A.L.,Y.A., J.D. and C.P.V. co-wrote the manuscript with input from all coauthors. All authors discussed the results and commented on the manuscript.

## DECLARATION OF INTERESTS

The authors declare that they have no competing interests.

## Supplementary Information

**Figure S1.**
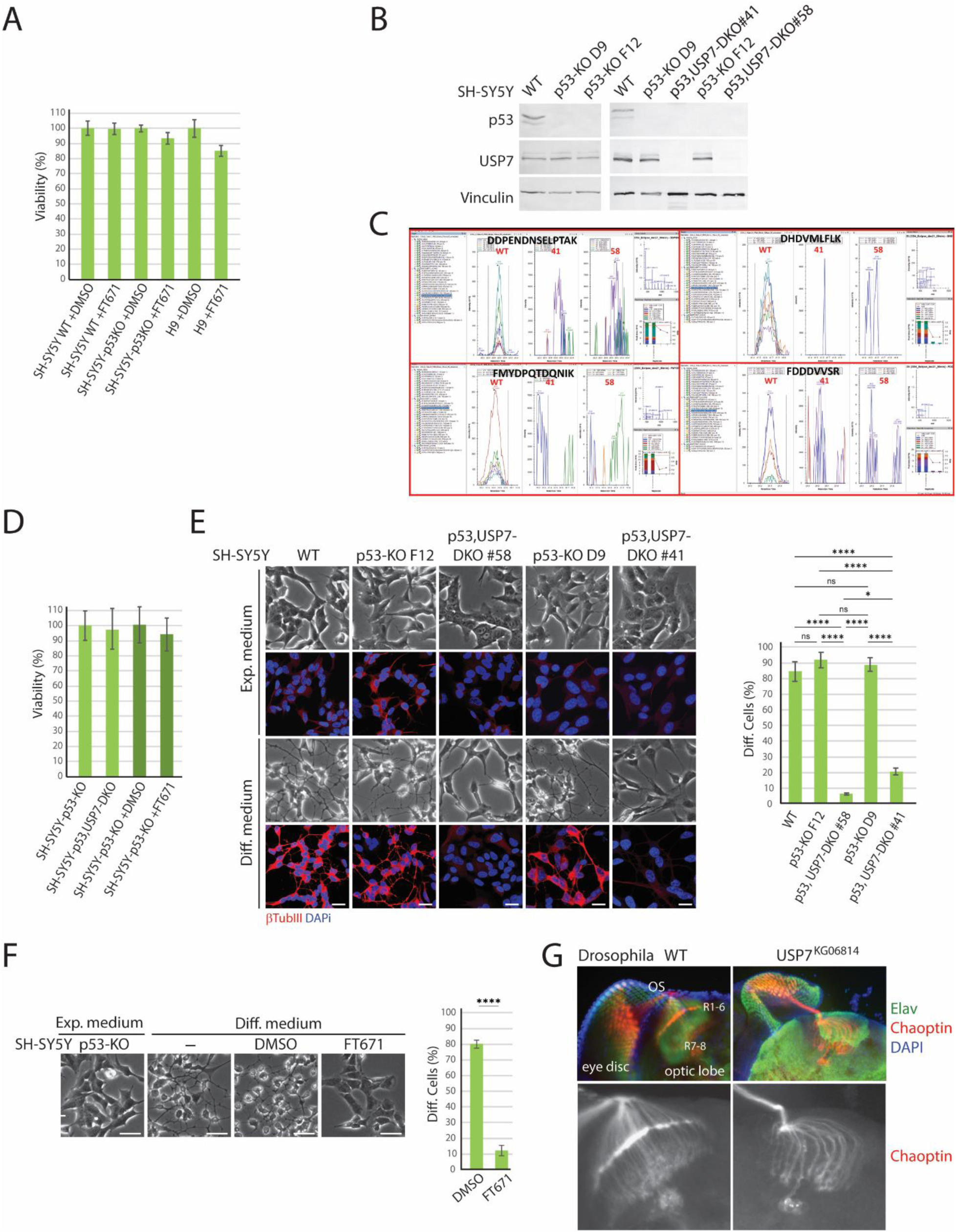
USP7 activity is crucial for neuronal differentiation of SH-SY5Y cell, related to. Figure 1 (A) Viability of SH-SY5Y and H9 cells in expansion medium (+/- FT671) as determined with the CellTiter®-Glo 2.0 assay. Means and SDs were derived from 3 biological replicates. (B) Immunoblot analysis of SH-SY5Y wt, and of 2 independent p53-KO clones and 2 independent p53,USP7-DKO clones. Vinculin serves as a loading control. p53,USP7-DKO#41 was derived from p53-KO D9; p53, USP7-DKO#58 was derived from p53-KO F12. (C) Targeted PRM MS analysis of 4 representative tryptic USP7 peptides in cell lysates of WT, #41 and #58 revealed their absence in USP7-KO lines. (D) Viability of p53-KO and p53, USP7-DKO SH-SY5Y cells in differentiation medium, as determined with the CellTiter®-Glo 2.0 assay. Means and SDs were derived from 3 biological replicates. (E) Effect of USP7 deletion on neuronal differentiation of SH-SY5Y cells. IF images of wt, and of 2 independent p53-KO clones and 2 independent p53,USP7-DKO clones, cultured in expansion or differentiation medium. Analysis as described for Figure 1C. See Data S2 for quantification. ****: p-value < 0.0001, *: p-value < 0.05, ns: not significant. (F) Effect of USP7i on neuronal differentiation of p53-KO F12 cells. (G) IF confocal sections of the developing visual system in wt and USP7^KG^ mutant *Drosophila* 3^rd^ instar larvae. Antibodies directed against Elav (Green) mark neurons and Chaoptin (Red) identify the photoreceptors and their axons (R1-8). OS: optic stalk. DNA was visualized by DAPI staining (blue).

**Figure S2.**
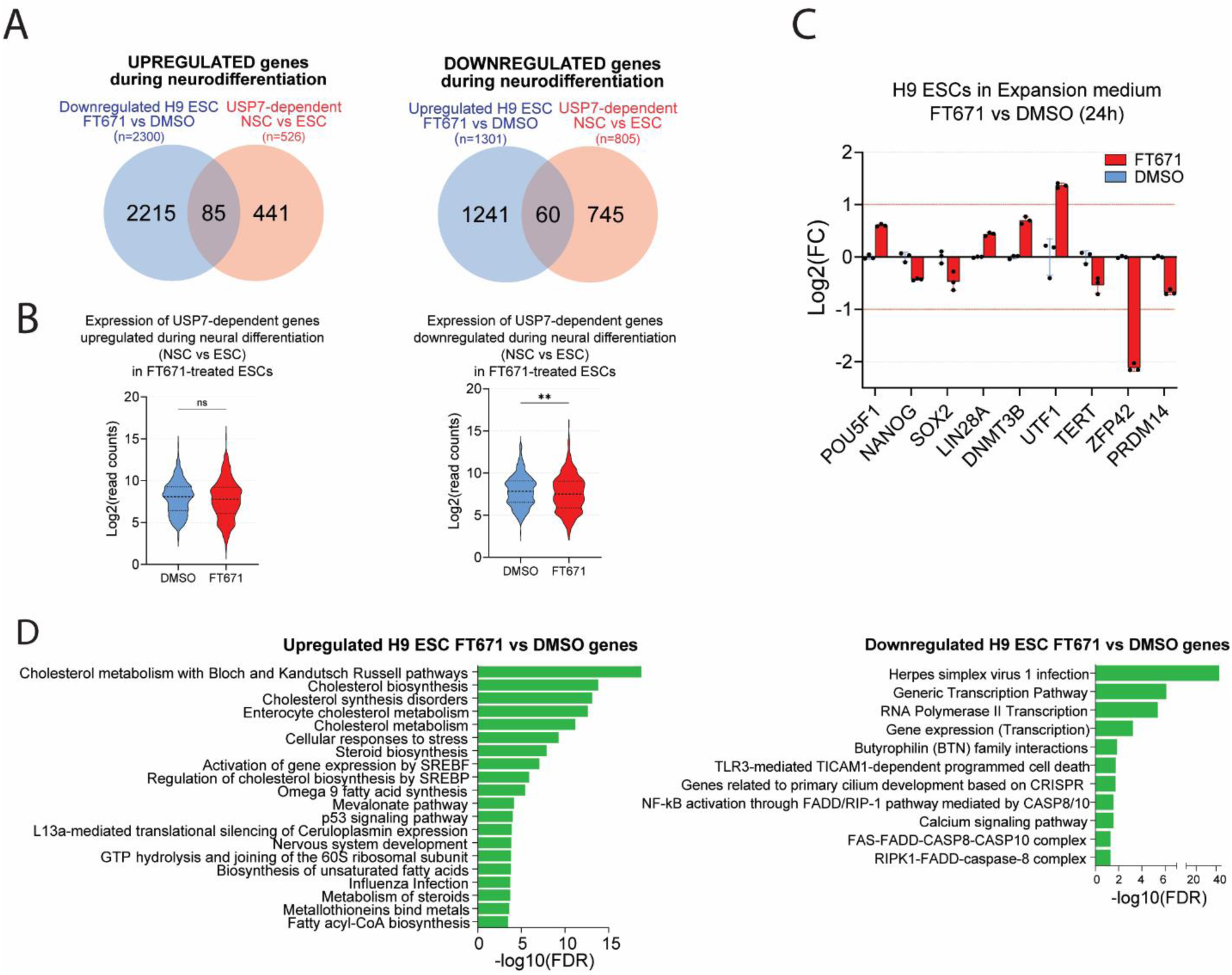
Transcriptional profiling of USP7i in H9 ESC cells grown in expansion medium, related to Figure 2 (A) Venn diagrams showing the overlap of DEGs in H9 ESCs grown in expansion medium in response to USP7i (24 hrs) and USP7 dependent DEGs in NSCs versus ESCs. (B) Violin plots of USP7 dependent DEGs in NSCs versus ESCs in ESCs grown in expansion medium (+/-FT671). Fold change reflects gene expression in the absence or presence of FT671. Means were derived from 3 biological replicates per condition. N.s. not significant, **: p-value < 0.01. (C) Analysis of pluripotency marker genes mRNA expression of H9 ESCs grown in expansion medium (+/-FT671). Values were based on RNAseq data. Means and standard deviations (SD) were derived from three biological replicates. (D) GO term analysis (KEGG) of DEGs in ESCs grown in expansion medium (+/-FT671).

**Figure S3.**
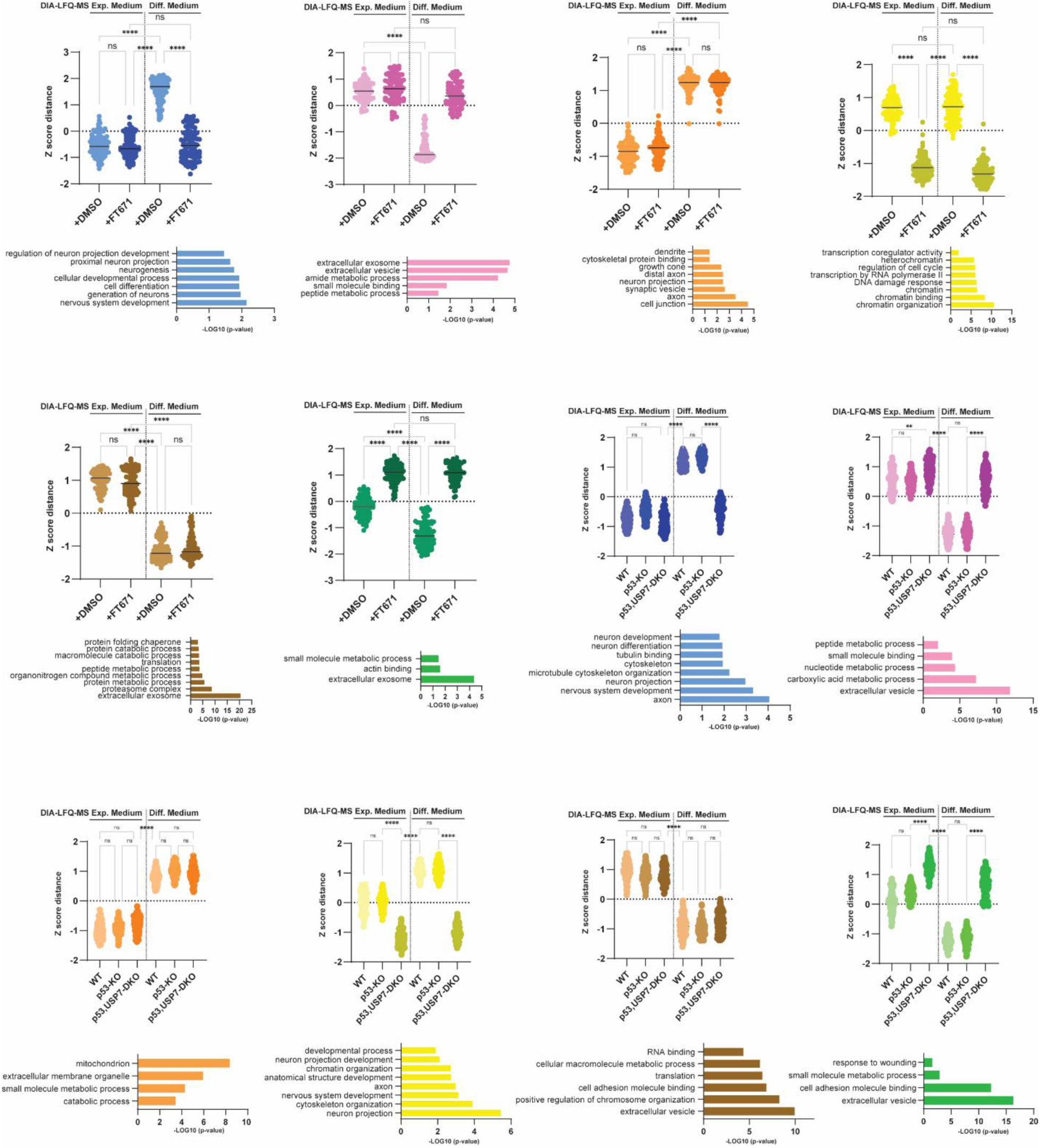
DIA-MS analysis of the USP7 network during neuronal differentiation, related to Figure 3 GO analysis of the top 100 proteins in wt SH-SY5Y cells that fit 6 different profiles following culture for 10 days in expansion- or in differentiation medium in the presence or absence of FT671, and a similar comparison of wt, p53-KO and p53,USP7-DKO SH-SY5Y cells (3 biological replicates per condition). For the complete data set see Data S4 and DataS5.

**Figure S4.**
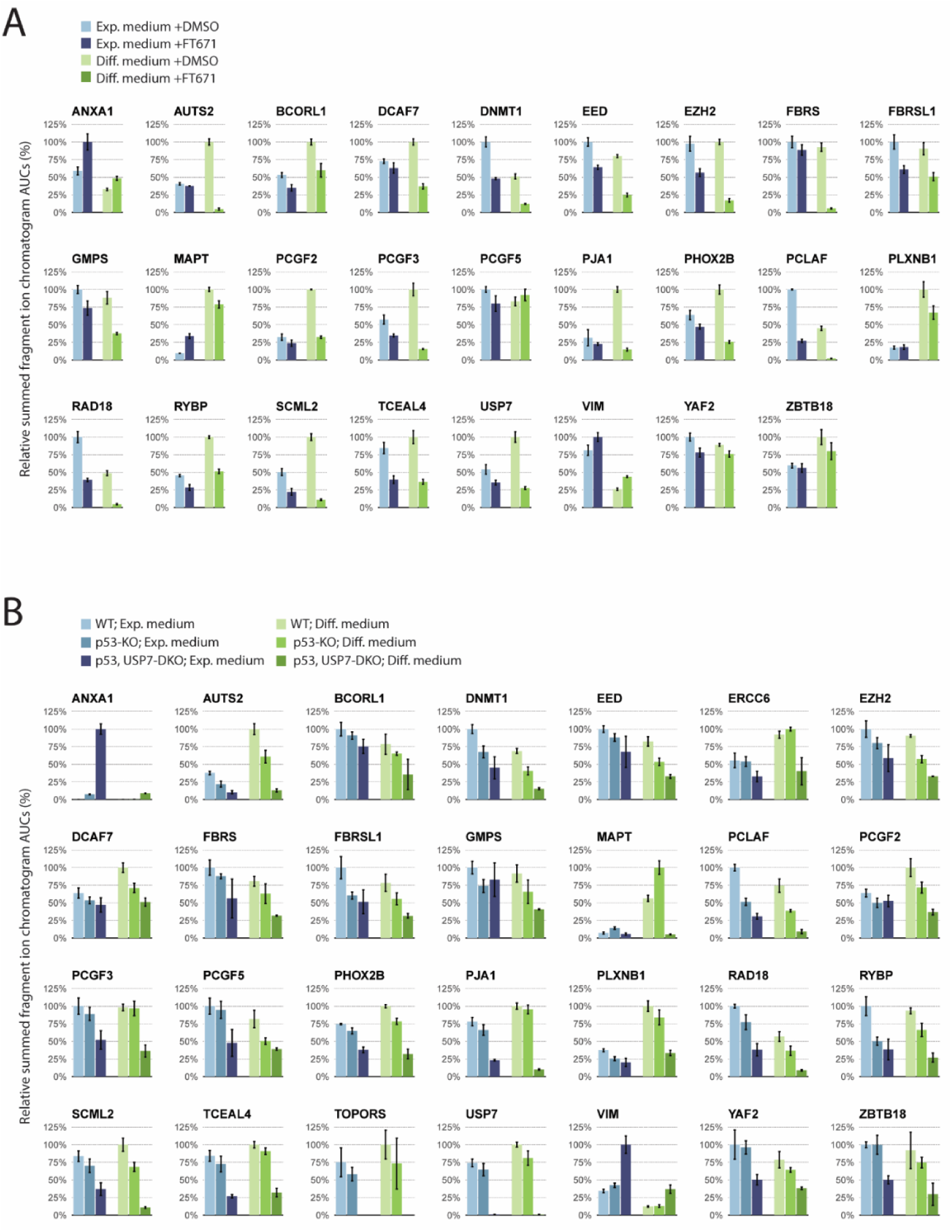
Quantitative PRM-MS analysis of the USP7 regulatory network, related to Figure 3 (A) PRM analysis of relative protein abundance in wt SH-SY5Y cells grown for 10 days in expansion- or in differentiation medium in the presence or absence of FT671 (3 biological replicates per condition). Chromatographic areas under the curve (AUCs) of peptides were summed to yield their respective protein quantities, which were subsequently averaged over three replicates and are shown as a percentage (± SD) of the largest resulting mean. For the complete data set see Data S7. (B) PRM analysis of relative protein abundance in wt, p53-KO and p53,USP7-DKO SH-SY5Y cells grown for 10 days in expansion- or in differentiation medium (3 biological replicates per condition). For the complete data set see Data S7.

**Figure S5.**
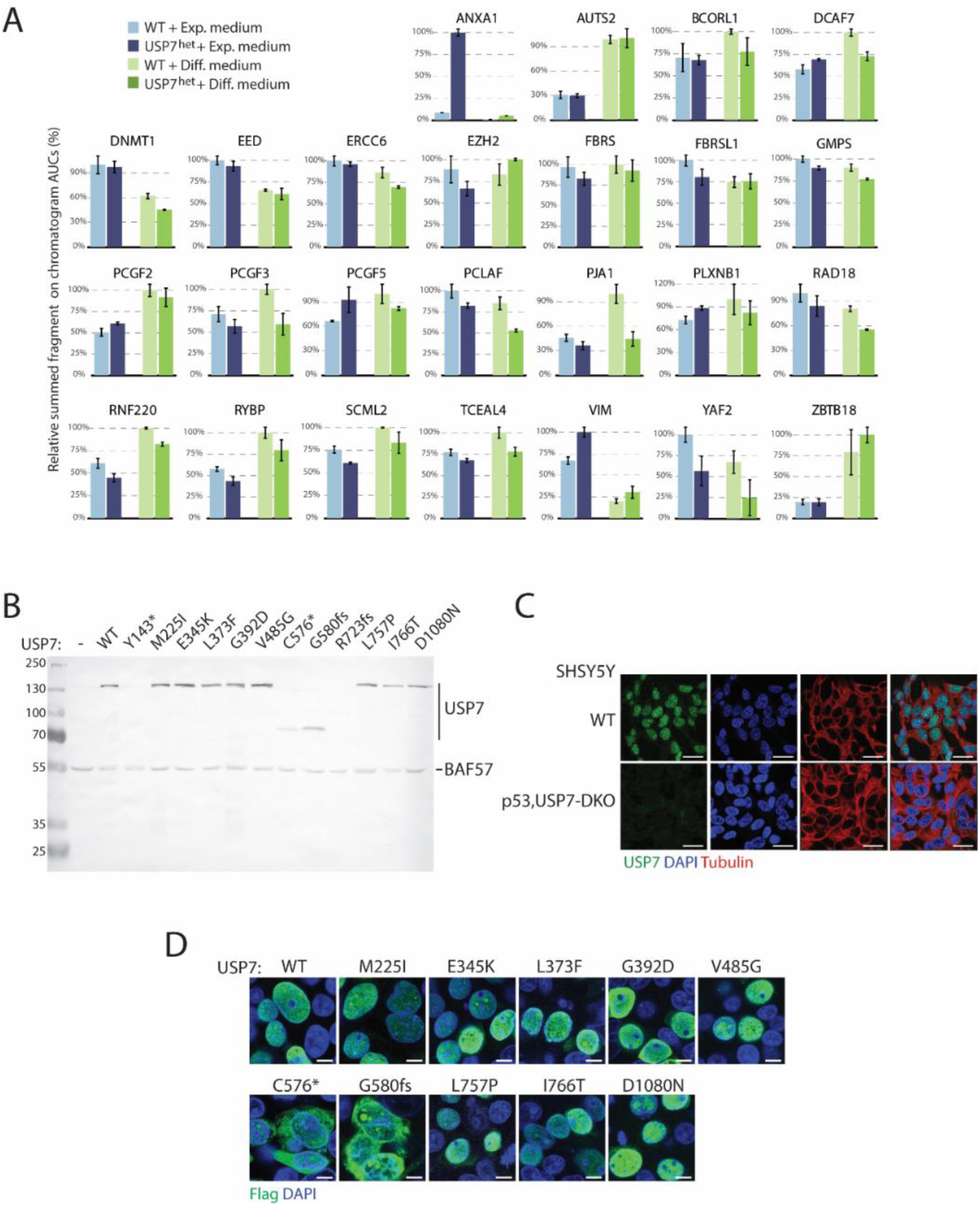
PRM-MS analysis of the impact of USP7 heterozygosity and expression of USP7 mutants associated with Hao-Fountain syndrome, related to Figure 4 (A) PRM analysis of relative protein abundance in wt and USP7^het^ SH-SY5Y cells grown for 10 days in expansion- or in differentiation medium (3 biological replicates per condition). Chromatographic areas under the curve (AUCs) of peptides were summed to yield their respective protein quantities, which were subsequently averaged over three replicates and are shown as a percentage (± SD) of the largest resulting mean. For the complete data set see Data S10. (B) Immunoblot analysis of USP7-KO HEK293T cells co-transfected with equal amounts of vectors expressing Flag-tagged wt mutant USP7. BAF57 serves as a loading control. (C) IF of endogenous USP7 using anti-USP7 antibodies in wt and p53,USP7-DKO cells. βTubIII was visualized by IF. DNA was visualized by DAPI staining (blue). Scale bar represents 25 μm. (D) IF analysis of Flag-tagged wt and mutant forms of USP7 expressed in USP7-KO HEK293T. Scale bar represents 10 μm.

**Figure S6.**
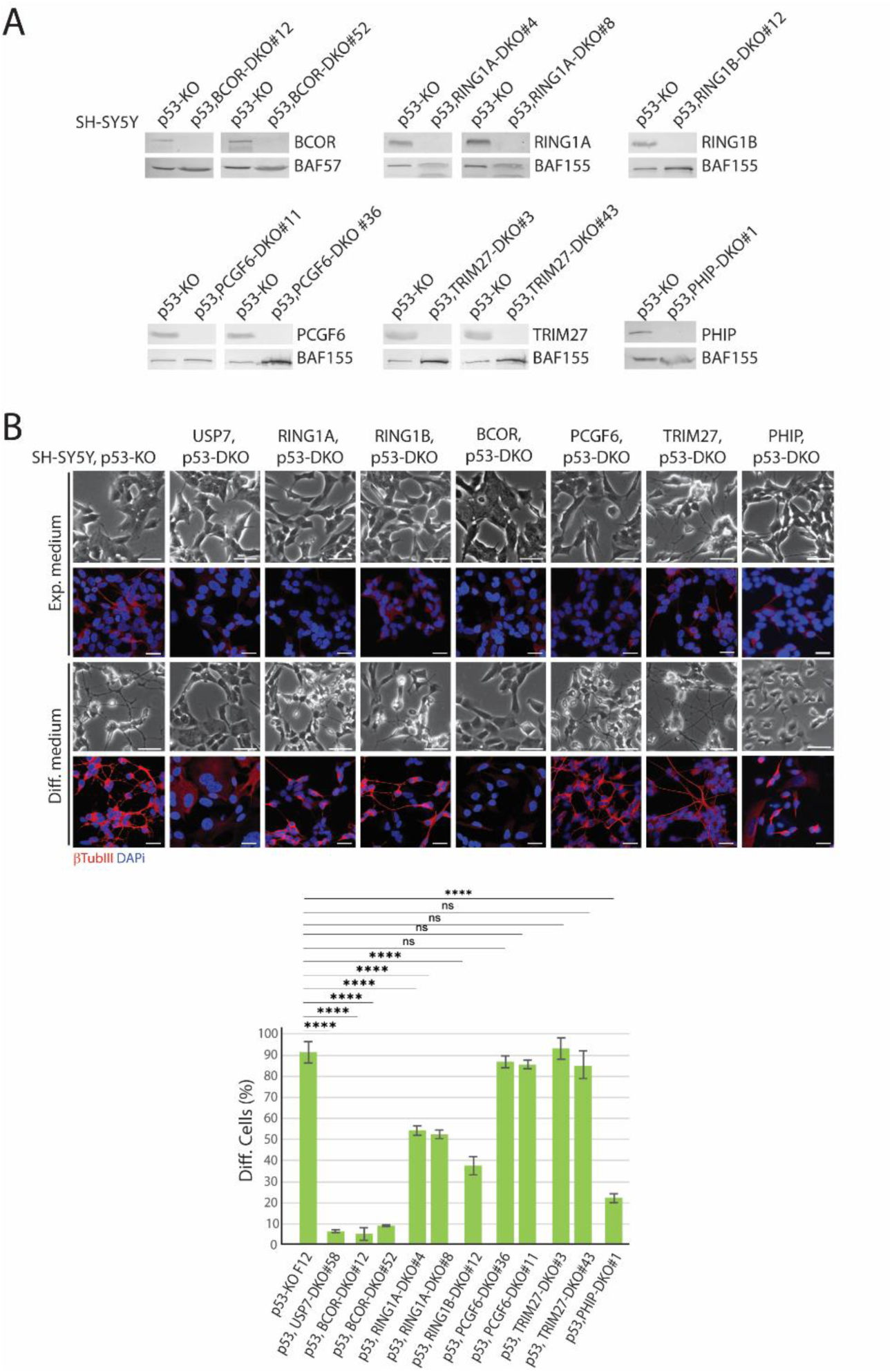
Effect of deletion of RING1A, RING1B, BCOR, PCGF6, TRIM27 or PHIP on neuronal differentiation of SH-SY5Y cells, related to Figure 5 (A) Immunoblot analysis of p53-KO SH-SY5Y cells and the derived clones in which either RING1A, RING1B, BCOR, PCGF6, TRIM27 or PHIP was deleted. BAF155 and BAF57 serve as loading controls. (B) Effect of deletion of RING1A, RING1B, BCOR, PCGF6, TRIM27 or PHIP on neuronal differentiation of p53-KO SH-SY5Y cells. The indicated DKO clones were cultured in expansion or differentiation medium for 10 days. The expression of βTubIII was determined by indirect IF. DNA was visualized by DAPI staining (blue). Scale bar represents 50 μm. ****: p-value < 0.0001, ns: not significant. Scale bar represents 50 μm. See Data S2 for quantification.

## Legend to Table S1

Table with critical materials.

Legends to Data S1 to S10 Data S1. (Separate excel file).

RNA-seq data of H9 ESCs grown in expansion or differentiation medium either in the presence or absence of FT671.

Data S2. (Separate excel file).

Percentage of differentiated cells, defined as having an extension from the cell body that is at least 2-times the length of the cell body of: i3N iPSCs, grown in the presence or absence of FT671; SH- SY5Y wt cells (+/-FT671); p53-KO (+/-FT671); p53,USP7-DKO; p53,RING1A-DKO; p53,RING1B-DKO; p53,BCOR-DKO; p53,PCGF6-DKO;p53,PHIP-DKO; p53,TRIM27-DKO; and USP7^het^ SH-SY5Y cells.

Data S3. (Separate excel file).

RNA-seq data of SH-SY5Y cells grown in expansion or differentiation medium: wt (+/-FT671); p53-KO; p53,USP7-DKO; p53,BCOR-DKO SH-SY5Y cells.

Data S4. (Separate excel file).

DIA-MS data from wt SHSY-5Y cells grown in expansion or in differentiation medium in the presence or absence of FT671.

Data S5. (Separate excel file).

DIA-MS data from wt, p53-KO and p53,USP7-DKO SH-SY5Y cells grown in expansion or in differentiation medium.

Data S6. (Separate excel file).

IP-MS data for proteins associated with endogenous USP7 purified from SH-SY5Y cells.

Data S7. (Separate excel file).

Peptide data from PRM analysis of relative protein abundance in wt (+/- FT671), p53-KO and p53,USP7-DKO SH-SY5Y cells grown in expansion- or in differentiation medium.

Data S8. (Separate excel file).

RNA-seq data of wt and USP7^het^ SH-SY5Y cells cultured in expansion or differentiation medium.

Data S9. (Separate excel file).

DIA-MS data from wt and USP7^het^ cells grown in either expansion or in differentiation medium.

Data S10. (Separate excel file).

Peptide data from PRM analysis of PRM-MS analysis of relative protein abundance in wt and USP7^het^ SH-SY5Y cells grown in expansion or in differentiation medium.

## Notes

### Competing Interest Statement

The authors have declared no competing interest.

### Summary of Updates

New results have been included in Figures 5A, 5B, S1, S2 and S6 5A and S6

